# Nucleocytoplasmic shuttling of the herpesvirus tegument protein UL47_TEG5 is required for its functional role in horizontal transmission in chickens

**DOI:** 10.64898/2026.09.08.750090

**Authors:** Hafiz Sohaib Zafar, Keith W. Jarosinski

## Abstract

Marek’s disease virus (MDV) UL47 tegument protein 5 (UL47_TEG5) is a major tegument protein essential for viral replication and horizontal transmission in chickens. The protein undergoes dynamic nucleocytoplasmic shuttling, likely regulated by host and viral factors, including post-translational modifications (PTMs). PTMs, in particular phosphorylation, on homologous herpesviral UL47_TEG5 proteins regulate nucleocytoplasmic shuttling through nuclear localization (NLS) and export (NES) motifs. Mass spectrometry-based proteomics of feather follicle epithelial skin from chickens infected with MDV recently showed that MDV UL47_TEG5 is phosphorylated. *In silico* analysis identified putative NLS and NES motifs near or within these phosphorylation sites. To test whether these motifs act as predicted, we systematically deleted the predicted NLS and NES motifs and assessed subcellular localization in transient transfection assays. Individual NLS deletions had minimal effect on nuclear localization, whereas NES deletions caused cytoplasmic accumulation and abolished nucleocytoplasmic shuttling. Next, we generated recombinant MDV lacking UL47_TEG5 (vΔ47) or the NES2 motif (aa 664-678) of UL47_TEG5 (v47ΔNES2) using two-step Red recombination. In experimentally infected chickens, replication and MD incidence were not different from those of wild-type MDV. However, in contact chickens, only wild-type MDV transmitted between birds, whereas both vΔ47 and v47ΔNES2 failed to transmit horizontally. These results demonstrate that nucleocytoplasmic shuttling of UL47_TEG5 is essential for horizontal transmission. In addition, our studies suggest that UL47_TEG5 expression, or lack thereof, in cells may be related to protein stability. Our findings link the nucleocytoplasmic shuttling of UL47_TEG5 to its essential function during natural infection and suggest this tegument protein may be a promising target for next-generation MDV vaccines.

**AUTHOR SUMMARY:** Marek’s disease virus (MDV) causes a highly contagious cancer in chickens and spreads efficiently from bird to bird. The viral protein UL47_TEG5 is essential for this horizontal transmission. We discovered that UL47_TEG5 constantly shuttles between the nucleus and cytoplasm, a process controlled by specific export signals (NES). Using molecular techniques, we deleted either NES motif, which trapped the protein in the cytoplasm and prevented shuttling. When we removed one key export signal (NES2) from the virus and infected chickens, the mutant virus replicated but completely failed to transmit to new birds. Our results show that UL47_TEG5’s ability to move between the nucleus and cytoplasm is critical for MDV transmission in chickens, likely through its function in virion assembly. These findings show that nucleocytoplasmic shuttling of UL47_TEG5 is mediated through a non-canonical mechanism and highlight UL47_TEG5 as a potential target for improved vaccines to better control this economically important disease.

## INTRODUCTION

Nucleocytoplasmic shuttling is fundamental to viral replication. Within cells, protein movement between the nucleus and cytoplasm regulates transcription, DNA replication, the cell cycle, and RNA transport, all of which are critical for both cellular function and viral replication. Viral proteins, such as human immunodeficiency virus (HIV) Rev, herpes simplex virus 1 (HSV-1) UL54_ICP27, and influenza NS2, were among the first proteins shown to undergo efficient nucleocytoplasmic shuttling [1–3]. Mislocalization or impaired shuttling disrupts virion assembly and ultimately affects infectious virion production [4, 5].

Herpesvirus tegument proteins play crucial roles in viral replication, gene regulation, and interactions with the host’s cellular machinery. Many of these proteins are multifunctional and shuttle dynamically between cellular compartments at distinct stages of infection. UL47_TEG5, or VP13/14, is a conserved alphaherpesvirus tegument protein that shuttles between the nucleus and cytoplasm (nucleocytoplasmic shuttling) [6–10]. UL47_TEG5 homologs, including those found in HSV-1, bovine herpesvirus 1 (BoHV-1), varicella-zoster virus (VZV), equine herpesvirus 1 (EHV1), and pseudorabies virus (PRV), localize to the nucleus, suggesting a role within the nucleus of infected cells. HSV-1 UL47_TEG5 has been shown to shuttle between the nucleus and cytoplasm [11] and to bind RNA through an N-terminal arginine-rich region that functions as a nuclear localization signal (NLS) [8]. HSV-1 UL47_TEG5 encodes a CRM1-dependent nuclear export signal (NES) in its C-terminus, and export is sensitive to Leptomycin B (LMB), consistent with classical CRM1-mediated export [7]. HSV-1 UL47_TEG5 compartmentalization is mediated by the serine/threonine protein kinase US3 via phosphorylation at a specific N-terminal serine residue (Ser-77), promoting its nuclear localization [12]. Disrupting phosphorylation through mutagenesis or inactivation of US3 kinase activity results in UL47_TEG5 accumulation at the nuclear periphery, impairing its nuclear localization. This localization leads to defects in viral replication and pathogenesis, underlying the biological significance of UL47_TEG5 localization. Similarly, US3-mediated phosphorylation of duck enteritis virus (DEV) UL47_TEG5 modulates the nucleocytoplasmic distribution in cells [13]. These findings suggest that UL47_TEG5 localization may be controlled by conditional switches, such as infection stage, binding partners, and phosphorylation status, rather than by a classical nuclear import/export pathway. Despite its conservation in alphaherpesviruses, the functional characterization and mechanism regulating UL47_TEG5 localization remain poorly understood in the host.

Gallid alphaherpesvirus 2 (species, *Mardivirus gallidalpha2*) or Marek’s disease virus (MDV) provides a unique model in which the functional relevance of viral protein localization and its role can be studied in a natural host system [14]. MDV is an alphaherpesvirus and causes Marek’s disease (MD) in chickens, characterized by immunosuppression, neurological symptoms, and rapid onset of T cell lymphomas. Natural infection begins with inhalation of infectious virus previously shed from chicken skin dander, and cytolytic infection initiates in B and T lymphocytes [15]. Infected cells then disseminate to lymphoid organs such as the spleen, bursa, and thymus, where MDV spreads through direct cell-to-cell contact. Infected immune cells also travel to the skin, where productive replication ensues in epithelial cells surrounding feather follicles, termed feather follicle epithelial (FFE) skin cells. In these cells, new infectious particles are produced and shed into the environment and horizontally transmitted to naïve chickens, a mode of dissemination similar to that of VZV in humans [16].

MDV UL47_TEG5 is required for horizontal transmission in chickens [17]. A recent study suggested its importance as related to its interaction with p32/C1QPB [18] and possibly linked to the regulation of *UL44* mRNA splicing and expression. *UL44* encodes glycoprotein C (gC), which is required for horizontal transmission of MDV in chickens [19, 20]. Studies comparing expression of MDV UL47_TEG5 showed that the expression and localization of UL47_TEG5 are considerably different between replication in cell culture and in the epithelial skin cells of infected chickens [21]. That is, MDV UL47_TEG5 is barely detectable and localizes primarily to the nucleus in cell culture, whereas in FFE skin cells, it is abundantly expressed and localizes to both the nucleus and the cytoplasm during the productive MDV replication cycle. This differential expression and localization of MDV UL47_TEG5 correlates with the cell-associated vs cell-free stages of MDV infection, suggesting UL47_TEG5 subcellular trafficking plays a functional role in the assembly and release of enveloped cell-free virus (CFV).

Here, we defined the subcellular trafficking of MDV UL47_TEG5 using a functional motif-mapping strategy that combines *in silico* prediction tools, quantitative cell-based localization assays, structural modeling, and an *in vivo* natural infection model in chickens. We identified putative NLS and NES motifs and demonstrated that the subcellular distribution of UL47_TEG5 is not governed by the classical import/export mechanisms. Inhibition of the classical export pathway with leptomycin B (LMB) and the importin α/β importation pathway with ivermectin (IVM) revealed that MDV UL47_TEG5 is insensitive to CRM1- and importin α/β-IV-mediated transport in cell culture, supporting regulation through a non-canonical pathway. Structural analysis using AlphaFold demonstrated that the NLS motifs are exposed and located in flexible regions, while NES motifs are embedded inside the core structure of the protein. The deletion of NES motifs did not disrupt global protein folding. Finally, we tested the role of UL47_TEG5 nucleocytoplasmic shuttling in chickens using a recombinant (r) virus (vUL47ΔNES) in which UL47_TEG5 could not translocate to the nucleus and failed to transmit, similar to a virus lacking UL47_TEG5 (vΔUL47). Together, these findings link UL47_TEG5 subcellular localization, structure, and biological function, providing a framework for understanding how the non-canonical subcellular trafficking of a herpesvirus tegument protein contributes to defective viral transmission in epithelial cells.

## RESULTS

### *In silico* predictions of NLS and NES motifs in MDV UL47_TEG5

Former studies in our laboratory showed that the localization of MDV UL47_TEG5 differed between cell-associated replication in cell culture and epithelial skin cells in chickens, where cell-free virus is produced [21, 22]. Chuard *et al.* [17] showed that MDV UL47_TEG5 was required for horizontal transmission in chickens, linking UL47_TEG5 expression and localization to cell-free virus production. Here, we aimed to determine the importance of UL47_TEG5 subcellular localization for its role in horizontal transmission. To identify putative nuclear localization (NLS) and nuclear export (NES) motifs within the MDV UL47_TEG5 protein, we first performed *in silico* analysis using multiple prediction tools. Classical monopartite and bipartite NLS were predicted with NLS Mapper [23], NLStradamus [24], NLSdb [25], seqNLS [26], and MyHits [27], while NES were predicted using LocNES [28]. High-scoring putative motifs are shown in Figure 1. All NLS-predicting tools consistently highlighted multiple clusters of basic residues (arginine-rich) in the N-terminal region of UL47_TEG5, particularly within residues 132-149 (NLS1) and 197-217 (NLS2), which corresponded to the strong scores. Another putative NLS was identified as a bipartite NLS [29] comprising two small basic clusters (RNR…KRT), separated by 21 residues, located from residues 270-300 (NLS3). Two leucine/isoleucine-rich motifs were predicted through the NES predictor as candidate canonical CRM-1-dependent nuclear export signals within 418-432 (NES1) and 664-678 (NES2) residues. Together, these computational prediction tools provided the first evidence that MDV UL47_TEG5 contains multiple candidate NLS and NES motifs. These predictions guided the design of subsequent mutagenesis and localization assays to experimentally validate these motifs.

**Fig 1.**
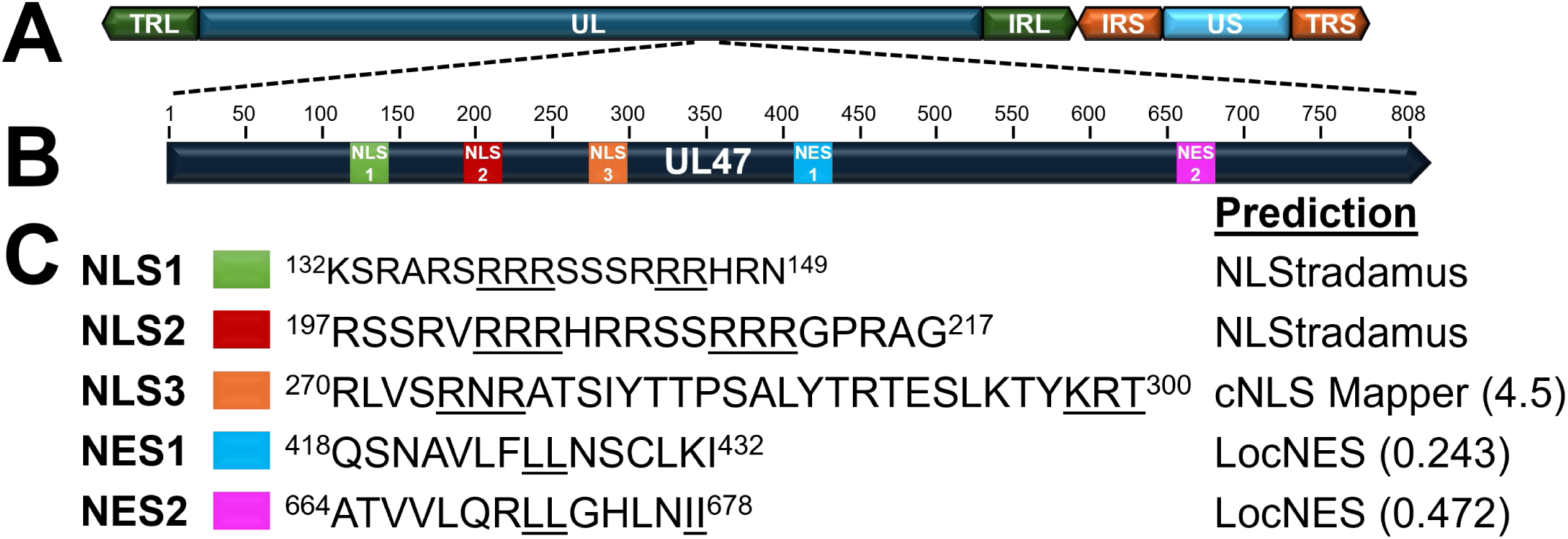
Schematic representation of MDV UL47_TEG5 NLS and NES motifs predicted in this report. (A) Schematic representation of the MDV genome, including the terminal repeat long (TRL) and short (TRS), internal repeat long (IRL) and short (IRS), and unique long (UL) and short (US) regions. (B) Full-length MDV UL47_TEG5 with annotation of predicted NLS1, NLS2, NLS3, NES1, and NES2 identified using *in silico* prediction tools. (C) The predicted NLS and NES amino acid sequences in MDV UL47_TEG5 are highlighted, and the prediction model used to derive the sequence and its score is shown. NLStradumus provides only the sequence, not a score.

### Truncation mapping reveals nuclear importation mediated by the C-terminal portion of MDV UL47_TEG5

Since MDV UL47_TEG5 is a large protein with many putative NLS or NES motifs, we started our investigation by generating an expression plasmid containing MDV UL47mRFP (pcUL47mRFP) cloned from previoulsy published recombinant (r) MDV, vUL47mRFP [30]. Then, we generated five in-frame truncations by deleting sequences 2-149 (Δ2-149), 150-300 (Δ150-300), 301-450 (Δ301-450), 451-600 (Δ451-600), and 600-808 (Δ600-808), as shown in Figure 2A. We used transient transfection assays and measured the nuclear-to-cytoplasmic ratio of the fluorescently tagged UL47_TEG5, as previously described in our laboratory [31]. Wild-type UL47mRFP was mostly nuclear. The two N-terminal truncations (Δ2-149 and Δ150-300) showed no change in N/C ratio with prominent nuclear localization, while the C-terminal truncations (Δ301-450, Δ451-600, and Δ600-808) shifted dramatically toward cytoplasmic distribution (Fig. 2B). Quantitative analysis of truncated proteins localization showed significantly lower N/C ratios for C-terminal truncations (N/C <1), reflecting enhanced cytoplasmic localization, while full-length (UL47mRFP) and N-terminal truncations (Δ2-149 and Δ150-300) maintained N/C ratios of >3 (Fig. 2C). These data indicate that the multiple regions within UL47_TEG5 contribute to nucleocytoplasmic shuttling, with deletion of C-terminal half of the protein significantly altering N/C ratio. Interestingly, these regions encode putative NESs, which would typically reduce nuclear export and increase nuclear localization. However, we saw the opposite result.

**Fig 2.**
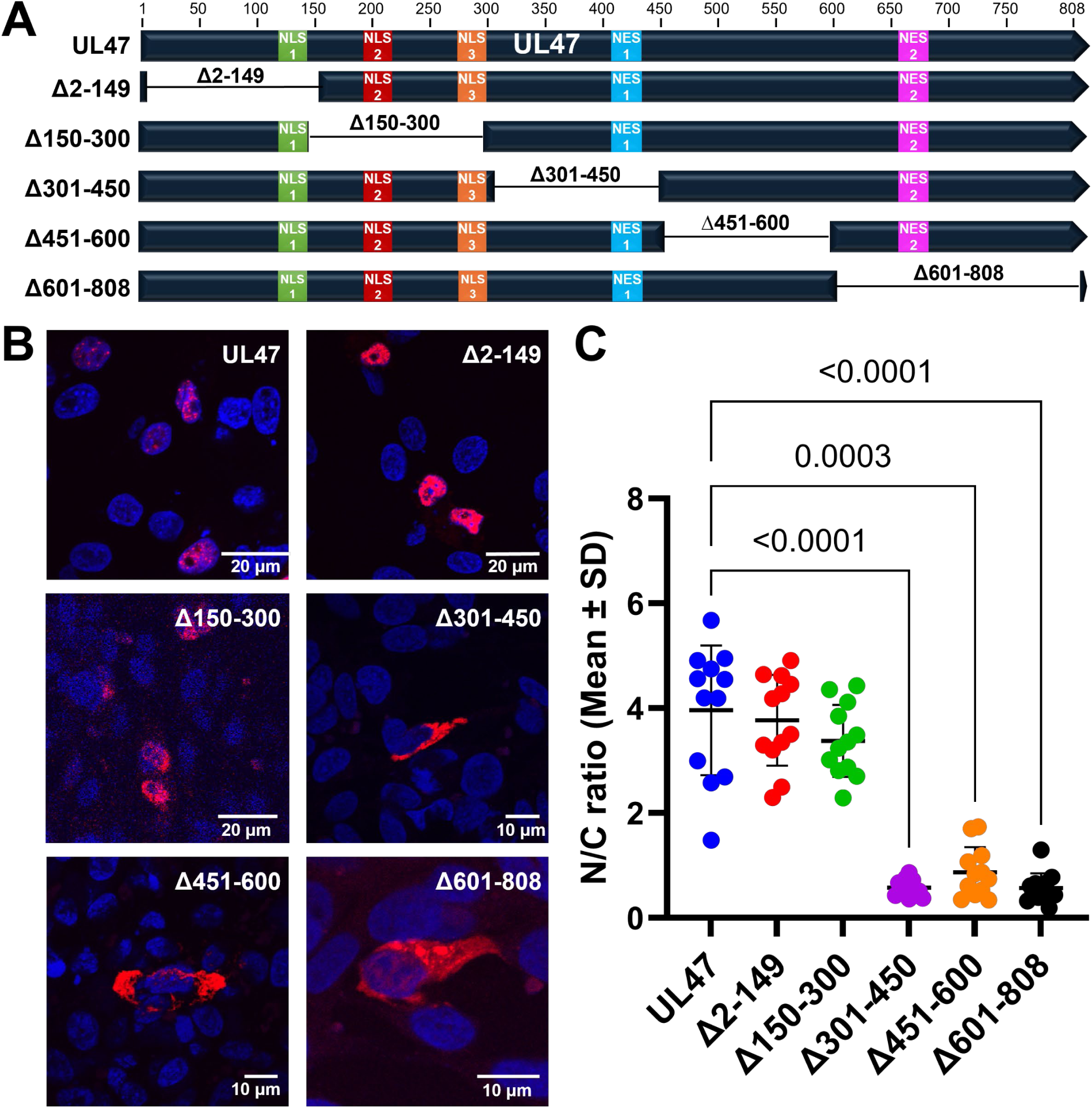
Subcellular localization of UL47_TEG5 truncation mutants. (A) Schematic representation of MDV UL47_TEG5 and truncation mutants generated in this report. (B) Representative images of UL47mRFP localization following transfection with pcUL47mRFP, pcΔ2-149, pcΔ150-200, pcΔ301-450, pcΔ451-600, and pcΔ601-808 using confocal microscopy. DF-1 cells were transfected with pcUL47mRFP (UL47) or each respective UL47mRFP-tagged mutant using Lipofectamine 2000. Cells were fixed with 4% PFA in PBS buffer at 24-36 h post-transfection and stained with Hoechst (DAPI) to visualize nuclei. Scale bars are 10 or 20µm. (C) Mean nuclear-to-cytoplasmic ratios (N/C ratios) were computed, and individual quantified cells (n= 12) are shown in a scatter dot plot. Data normality was assessed using the Shapiro-Wilk test. We used a Kruskal-Wallis test followed by Dunn’s multiple-comparison test to assess differences between groups. Significant differences between the UL47mRFP and truncation mutants are indicated by p-values. Data are presented as the mean ± standard deviation (SD). p > 0.05 (not significant), p < 0.05 (significant), p < 0.001 (highly significant).

### The effect of combined UL47_TEG5 NLS motif deletions on subcellular localization

It was interesting that deleting the N-terminal 300 aa of UL47_TEG5 did not alter its prominent nuclear localization, since this region typically encodes NLS motifs in other herpesviruses [6, 32]. Therefore, we further defined this region specifically to target predicted NLS motifs. To do this, we generated double and triple mutants (ΔNLS1+2, ΔNLS2+3, ΔNLS1+3, ΔNLS1+2+3) that retain single NLS motifs (Fig. 3A). In transient transfection assays, all combined deletion mutants retained predominant nuclear localization (Fig. 3B). However, there was a significant shift in N/C ratio (4.14 to 2.06) when all three NLS were deleted (ΔNLS1+2+3) compared to full-length UL47 (Fig. 3C). These findings suggest that the deletion of certain N-terminus regions shifts the equilibrium of UL47_TEG5 subcellular localization.

**Fig 3.**
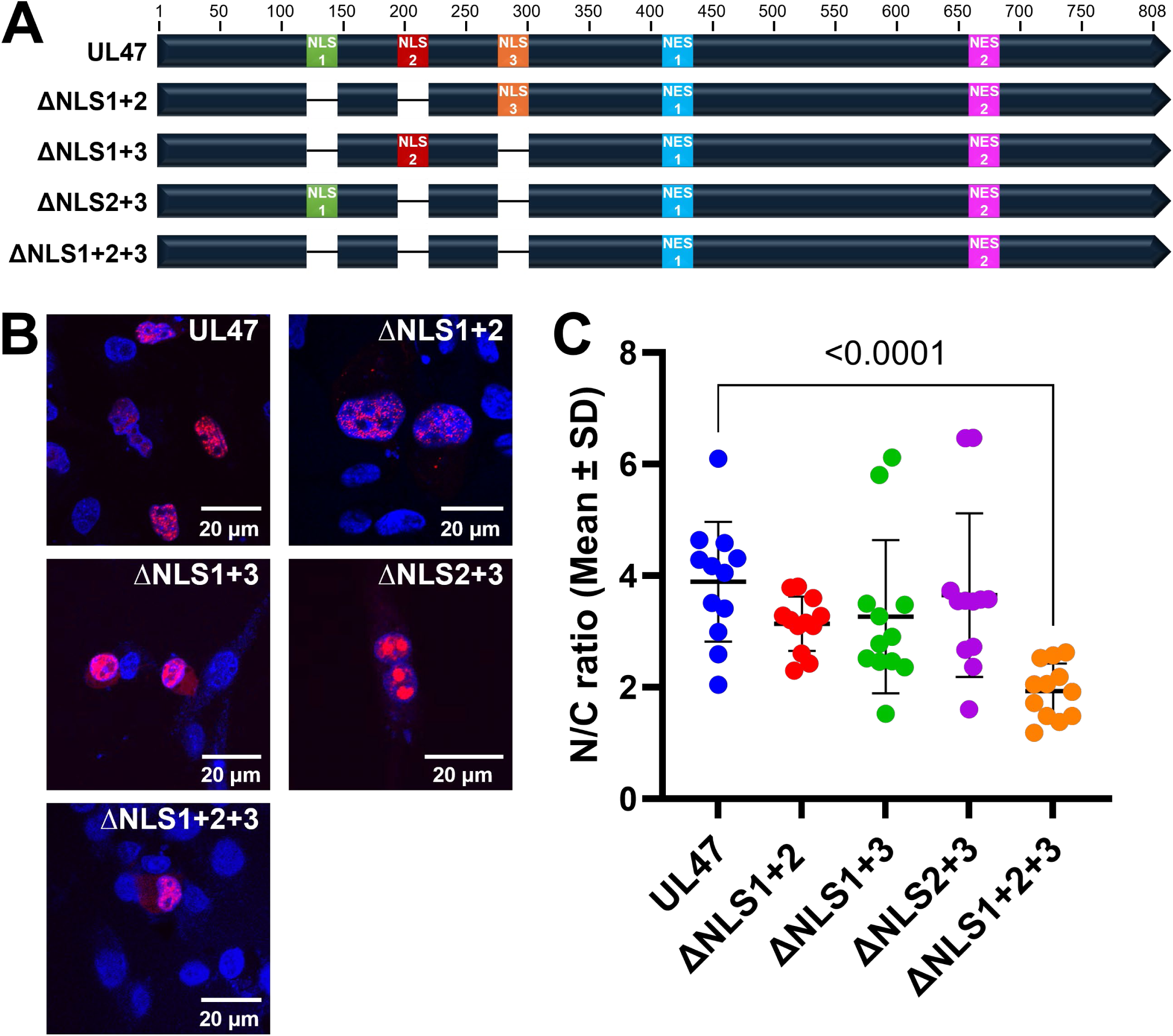
Subcellular localization of UL47_TEG5 double/triple mutants. (A) Schematic representation of MDV UL47 double/triple NLS_TEG5 mutants generated in this report. (B) Representative confocal images of UL47 double/triple mutations, pcΔNLS1+2, pcΔNLS1+3, pcΔNLS3+3, and pcΔNLS1+2+3. DF-1 cells were transfected with the pcUL47mRFP (UL47) or the respective mutant expression constructs using Lipofectamine 2000. Cells were fixed with 4% PFA at 24-36 h post-infection and stained with Hoechst 33342 to visualize nuclei. Scale bars are 20 µm. (C) Mean nuclear-to-cytoplasmic ratios (N/C ratios) were computed, and individual quantified cells (n= 12) are shown in a scatter dot plot. We assessed data normality using the Shapiro-Wilk test. We used the Kruskal-Wallis test followed by Dunn’s multiple-comparison test to assess differences between groups. Significant differences between the UL47 and double/triple deletion mutants are indicated by p-values. Data are presented as the mean ± SD.

### The effect of individual UL47_TEG5 NLS motif deletion on subcellular localization

To further assess whether UL47_TEG5 relies on a single classical nuclear localization signal for nuclear import, we examined the subcellular localization of UL47_TEG5 by generating individual deletions of each predicted NLS motif, that retain two predicted NLS (Fig. 4A). Deletion of individual NLS motifs (ΔNLS1, ΔNLS2, ΔNLS3) did not alter the nuclear localization of UL47_TEG5 (Fig. 4B). All NLS deletion mutants retained a predominant nuclear localization signal, indicating that no single NLS is strictly required for UL47_TEG5 nuclear import (Fig. 4C).

**Fig 4.**
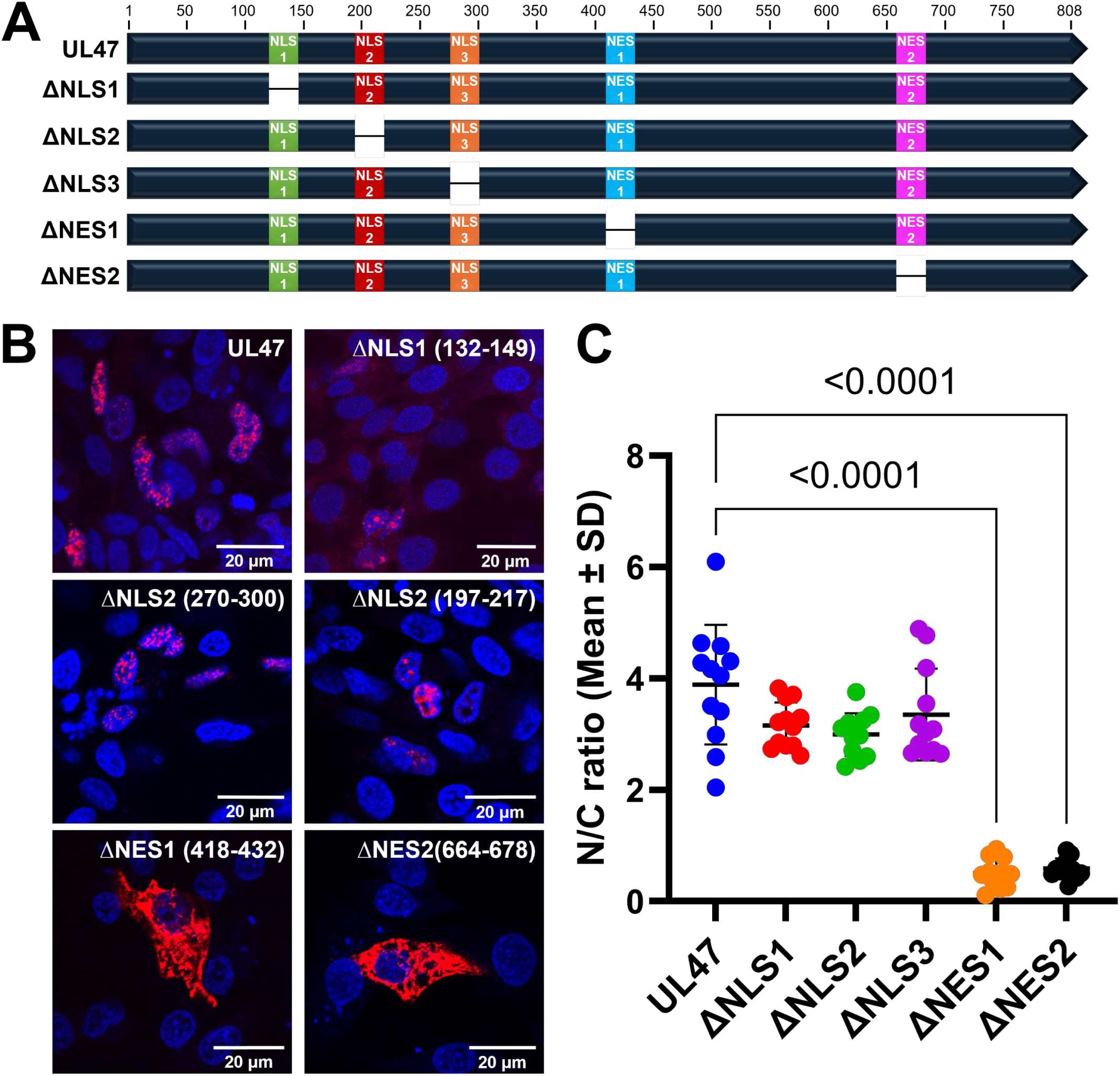
Subcellular localization of UL47_TEG5 individual NLS and NES mutants in DF-1 cells. (A) Schematic representation of MDV UL47_TEG5 individual deletion mutants generated in this report. (B) Representative confocal images of pcUL47mRFP (UL47) and individual NLS/NES mutations. DF-1 cells were transfected with the pcUL47mRFP (UL47) and respective mutant expression constructs (pcΔNLS1, pcΔNLS2, pcΔNLS3, pcΔNES1, and pcΔNES2) using Lipofectamine 2000. Cells were fixed with 4% PFA in PBS at 24-36 h post-infection and stained with Hoechst 33342 to visualize nuclei. Scale bars are 20µm. (C) Mean nuclear-to-cytoplasmic ratios (N/C ratios) were computed, and individual quantified cells (n= 12) are shown in a scatter dot plot. Data normality was assessed using the Shapiro-Wilk test. We used a Kruskal-Wallis test followed by Dunn’s multiple-comparison test to assess differences between groups. Significant differences between the UL47 and single deletion mutants are indicated by p-values. Data are presented as the mean ±SD.

#### Deletion of the NES motifs leads to unexpected cytoplasmic localization

Since deletion of putative NLS motifs individually or in combinations did not affect UL47_TEG5 nuclear localization, we next focused on the two predicted NES motifs (Fig. 4A). Each of the two putative leucine-rich NES motifs in pcUL47mRFP was deleted to generate ΔNES1 (Δ418-432) and ΔNES2 (Δ664-678) mutants. Localization assays revealed that deletion of either motif resulted in a clear shift toward cytoplasmic distribution compared to full-length UL47_TEG5 (Fig. 4B). This was also supported by quantification of the subcellular localization of these mutants, which showed a significantly reduced mean N/C ratio (4.05 to ∼0.5) (Fig. 4C). These results show that regions, NES1 (418-432) and NES2 (664-678), are directly involved in the subcellular localization of MDV UL47_TEG5 in cells. However, deletion of these regions reverses the effect on UL47_TEG5, shifting its localization in cells from ∼4-fold N/C to ∼0.5 N/C.

#### Effect of CRM-1 and importin α/β inhibition on UL47 subcellular localization

Under a canonical pathway, UL47_TEG5 nuclear localization would be expected to depend on importin α/β-mediated import via a defined NLS, while cytoplasmic localization would be mediated by CRM1-dependent export through a leucine-rich NES motif. Based on the localization effects of individual NLS and NES deletions, we hypothesized that UL47_TEG5 does not rely on a traditional nuclear import/export pathway but instead employs a non-canonical nucleocytoplasmic shuttling mechanism. To test this hypothesis, we examined the effects of pharmacological inhibition of nuclear import and export pathways on UL47_TEG5 localization. To do this, we transfected DF-1 cells with pcUL47mRFP (UL47), ΔNES1 (Δ418-432), or ΔNES2 (Δ664-678) expression constructs and treated them with 20 nM LMB for 6 h to inhibit CRM1-dependent export [33] or with 12.5 µM ivermectin (IVM) for 1.5 h to inhibit importin α/β-mediated nuclear import [34] prior to fixation and analysis by fluorescence assays. As a positive control for CRM1-dependent subcellular distribution, we used the EIAV-Rev EGFP construct previously shown to be mediated by CRM1-dependent export [35]. Without drug treatment, EIAV-Rev EGFP exhibited both nuclear and cytoplasmic localization (N/C = 1.37 ± 0.84) (Fig. 5). Upon LMB treatment, EIAV-Rev GFP redistributed to the nucleus exclusively (N/C = 12.19 ± 8.14), confirming effective inhibition of CRM1-dependent export, validating both the drug’s activity and the imaging conditions. IVM treatment reduced the N/C ratio from 1.37 ± 0.84 to 0.73 ± 0.81, though this was not significant. UL47_TEG5 remained predominantly nuclear (N/C = 4.90 ± 1.85) with no change following LMB or IVM treatments. Similarly, both ΔNES1 and ΔNES2 expression constructs retained their cytoplasmic distribution as observed during individual deletions (N/C < 1.0). These results show that MDV UL47_TEG5 localization is not mediated by canonical CRM1- or importin α/β-mediated pathways.

**Fig 5.**
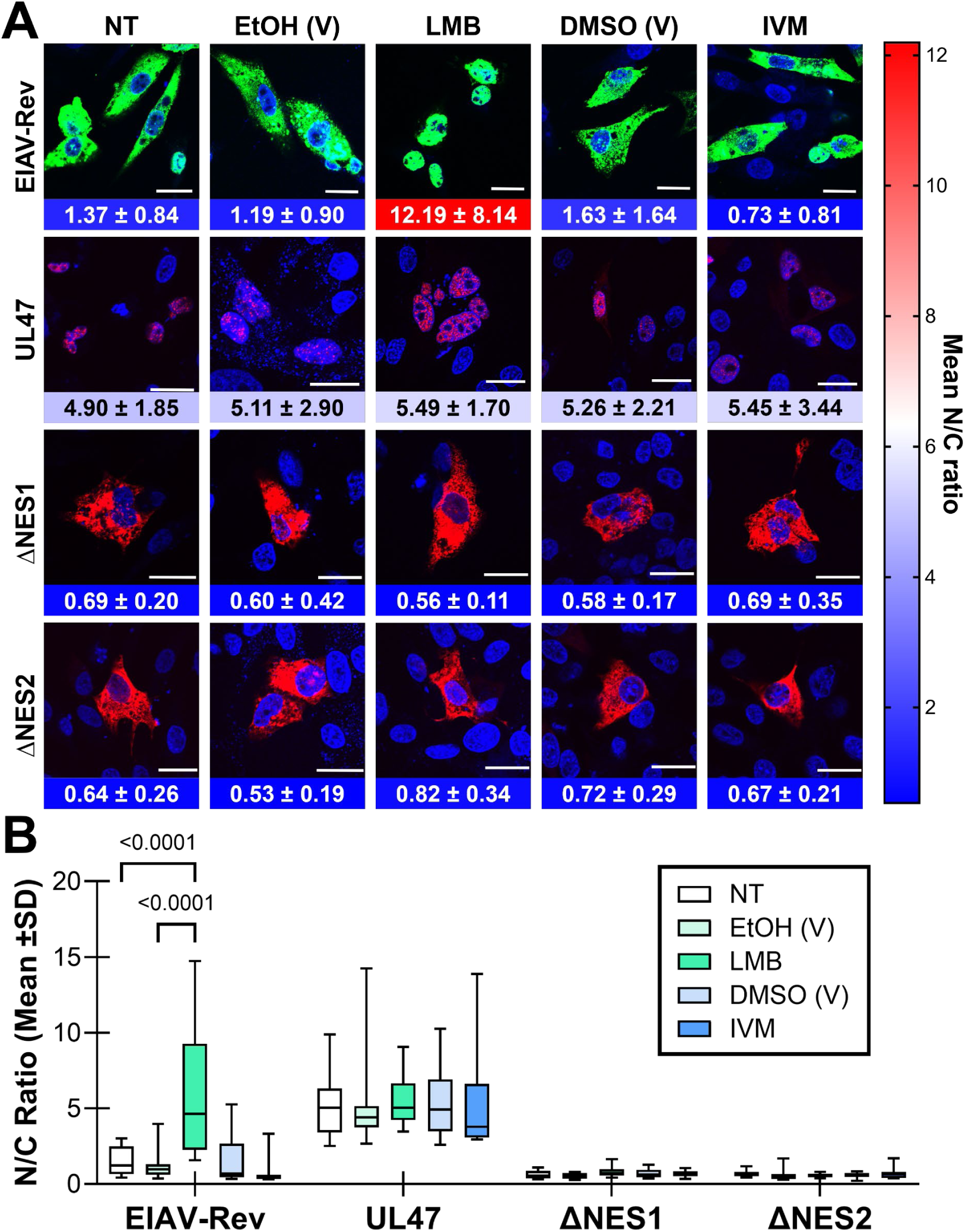
Localization of UL47, ΔNES1, and ΔNES2 following treatment with nuclear export and import inhibitors. (A) Representative confocal images showing localization of EIAV-Rev-EGFP (EIAV-Rev), MDV UL47, ΔNES1, and ΔNES2 following treatment with Leptomycin B (LB) or ivermectin (IVM). DF-1 cells were transfected using Lipofectamine 2000 with pcUL47mRFP (UL47), NES mutants, or EIAV-Rev as a positive control for CRM1-mediated export inhibition. After 24-36 hours post-infection, cells were treated with LMB (20 nM) or vehicle (ethanol) for 6 h to inhibit CRM1-mediated nuclear export, or IVM (12.5 μM) or vehicle (DMSO) for 1.5 h. After treatment, cells were fixed with 4% PFA, stained with Hoechst 33342, and processed for confocal microscopy. Scale bars are 20 µm. Mean nuclear-to-cytoplasmic ratios (N/C ratios) were computed, and individual quantified cells (n = 13) were represented as a heatmap. Data are presented as the mean ± SD. (B) We assessed data normality using the Shapiro-Wilk test. A two-way ANOVA followed by Tukey’s multiple-comparison test was performed to assess differences between groups in drug treatment. Significant differences between no treatment and drug treatment in the control (EIAV-Rev) and UL47_TEG5 mutants are indicated by p-values. NT= no treatment.

### Structural modeling of UL47_TEG5 using AlphaFold reveals little effect on structure by deleting NES motifs

To gain structural insight into the effect of deleting the predicted NLS or NES motifs of UL47_TEG5, we used AlphaFold2, implemented in the ColabFold pipeline [36, 37], to generate three-dimensional (3D) models of full-length UL47_TEG5 and its NES deletion mutants. First, we visualized the predicted structure of full-length UL47_TEG5 colored by predicted local distance difference test (pLDDT) scores, which provide residue-level confidence estimates for the AlphaFold predictions (Fig. 6A). The 3D model of full-length UL47_TEG5 showed a well-defined central core structure (aa 400-800) with a high pLDDT score indicative of a confidently predicted domain. In contrast, the N-terminal region exhibited lower pLDDT values, consistent with low confidence and disorder. All three putative NLS regions were mapped to this lower-confidence region, suggesting that these motifs may function within a flexible, disordered region of the protein. Individual predicted motifs were colored separately on the 3D model of UL47_TEG5, and the NES motifs are highlighted, demonstrating that these motifs are anchored within the structural core of UL47_TEG5 rather than exposed on the surface (Fig. 6B).

**Fig 6.**
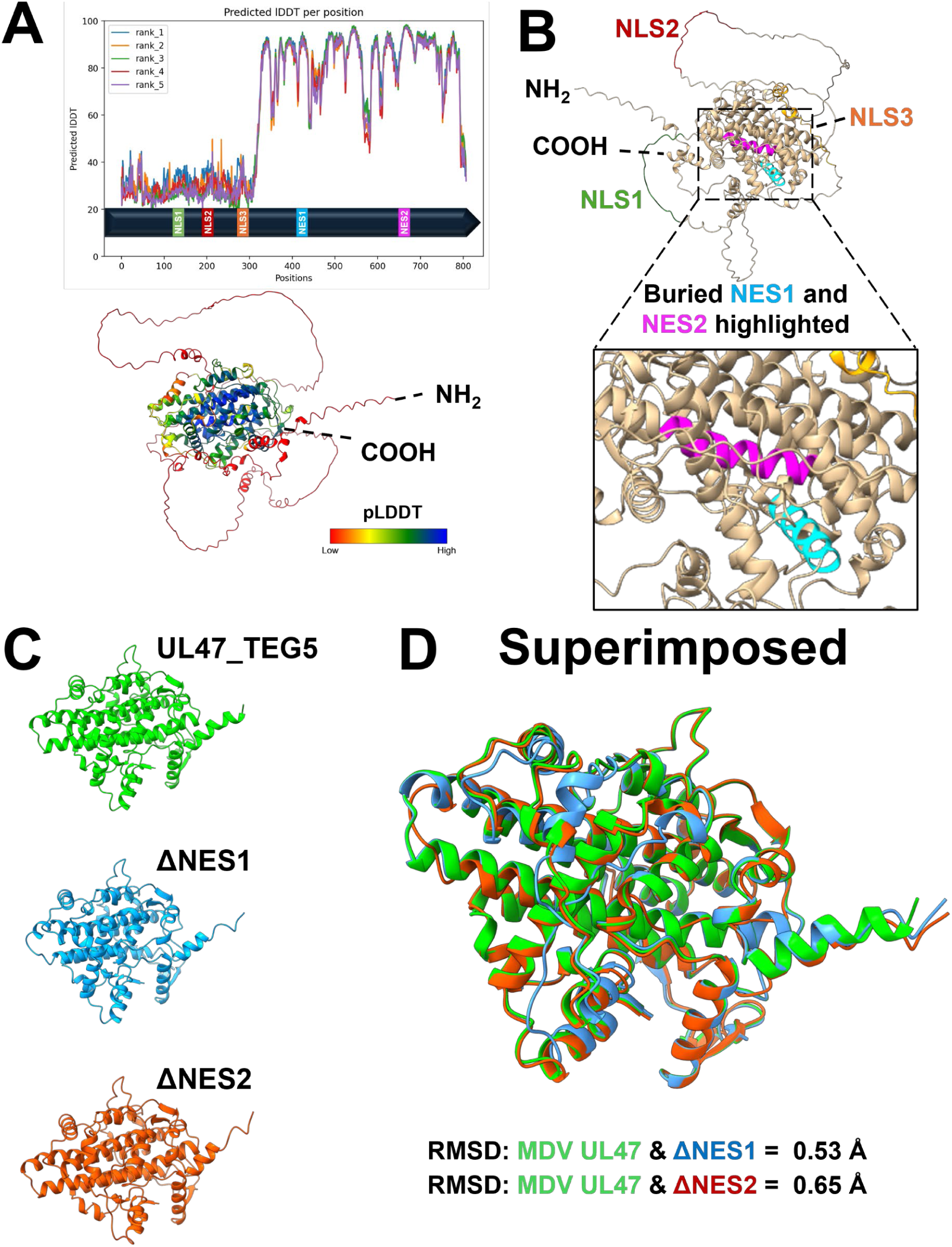
AlphaFold structural modeling of MDV UL47_TEG5 and ΔNES motif mutants. (A) Predicted structure of UL47_TEG5 (UniProt: Q9E6M8), colored by per-residue confidence (pLDDT), with high-confidence regions shown in cooler colors and low-confidence regions in warmer colors. (B) Putative NLS and NES motifs are highlighted in color on the structural model. The C-terminal region containing NES1 (aa 418-432) and NES2 (aa 668-675) motifs was focused on to show that it is embedded within the UL47_TEG5 core structure. (C) Individual view of UL47_TEG5, ΔNES1, and ΔNES2 core regions comprising residues with high pLDDT scores. (D) Structural superimposition of UL47_TEG5, ΔNES1, and ΔNES2 mutants, with root mean square deviation (RMSD) calculated across aligned residues. Low RMSD values across aligned residues indicate no global misfolding or structural perturbations.

To further assess the confidence of these regions, we examined predicted aligned error (PAE) plots generated for the top-ranked AlphaFold2 models. PAE analysis revealed low-error blocks in the central core region (aa 400-800), indicating high confidence in the results (Fig. S1A). In contrast, higher PAE values were observed between the core and terminal regions, indicating uncertainty in the orientation of this region (aa 1-400). The predicted NLS motifs are primarily located in the high-PAE and low-confidence regions, supporting the idea that these fragments are disordered, conformationally flexible, and may adopt multiple orientations. We did not observe any large-scale differences in the core domain when comparing PAE plots of full-length UL47_TEG5 and NES-deletion mutants, suggesting that removing NES regions does not globally destabilize UL47_TEG5 folding.

Next, we assessed whether deletion of the two NES regions affected UL47_TEG5 folding. We isolated the core domain with high confidence from AlphaFold2 and independently evaluated its structural quality. The low-confidence N-terminal region was removed prior to visualization of the core domain (Fig. 6C). Model quality was further evaluated using MolProbity-based stereochemical validation. Ramachandran analysis of all three UL47_TEG5 core domain structures showed that most residues were in favored regions, with few outliers. The overall Ramachandran z-score was consistent with a well-formed protein structure (Fig. S1B). We performed a direct structural comparison by superimposing the UL47_TEG5, ΔNES1 (Δ418-432), and ΔNES2 (Δ664-678) core models using ChimeraX’s matchmaker. The superimposed structure of the wild-type core and corresponding NES deletion mutants revealed low root mean square deviation (RMSD) values, indicating that the removal of NES motifs does not disrupt the global structure of the UL47_TEG5 core (Fig. 6D).

These structural findings provide important insights into interpreting the functional data observed in our localization assays. Deletion of NES motifs resulted in enhanced cytoplasmic distribution of UL47_TEG5, a phenotype that is counterintuitive under traditional CRM-1-dependent export. In contrast, structural analysis suggests that NES motifs contribute to UL47_TEG5 localization through nuclear retention or interactions with nuclear partners. Removing these motifs likely disrupts these interactions, leading to cytoplasmic redistribution while preserving overall protein folding. These structural understandings support a model in which UL47_TEG5 NES motifs play a central role in determining steady-state nucleocytoplasmic movement, and observed phenotypes in localization assays arise from altered trafficking or mislocalization of UL47_TEG5 rather than from global misfolding or structural instability.

### MDV UL47_TEG5 NES2 is required for nuclear localization of UL47_TEG5 during infection

NES2 is required for the nuclear localization of MDV UL47_TEG5 during transient transfection in DF-1 cells (Fig. 4). Next, we determined whether NES2 is required for UL47_TEG5 localization during infection by generating a mutant rMDV in which the NES2 region was deleted. Since MDV UL47eGFP is barely detectable in cell culture, we used a previously published recombinant (r)MDV in which MDV UL47eGFP is driven by the CMV promoter called vPcmvUL47eGFP [22]. This is clearly shown in Figure 7, where virtually no UL47eGFP can be detected in vUL47eGFP-infected chick embryo cells (CECs), while it can be clearly observed in vPcmv47eGFP-infected cells. To generate vPcmv47ΔNES2, aa 664-678 were deleted using two-step recombination (Fig. S2). All recombinant viruses were screened by restriction fragment length polymorphism (RFLP) analysis to confirm the integrity of the bacterial artificial chromosome (BAC) clones. Following reconstitution of BAC clones into virus stocks, confocal microscopy was used to examine the localization of UL47eGFP. A large shift from nuclear to cytoplasmic localization of UL47eGFP was observed comparing vPcm47ΔNES2 to vPcmvUL47eGFP (Fig. 7B). The N/C ratio of UL47eGFP changed from >3-fold to <0.5-fold between each virus, confirming NES2 (aa 664-678) is required for nuclear localization during replication in cell culture (Fig. 7C).

**Fig 7.**
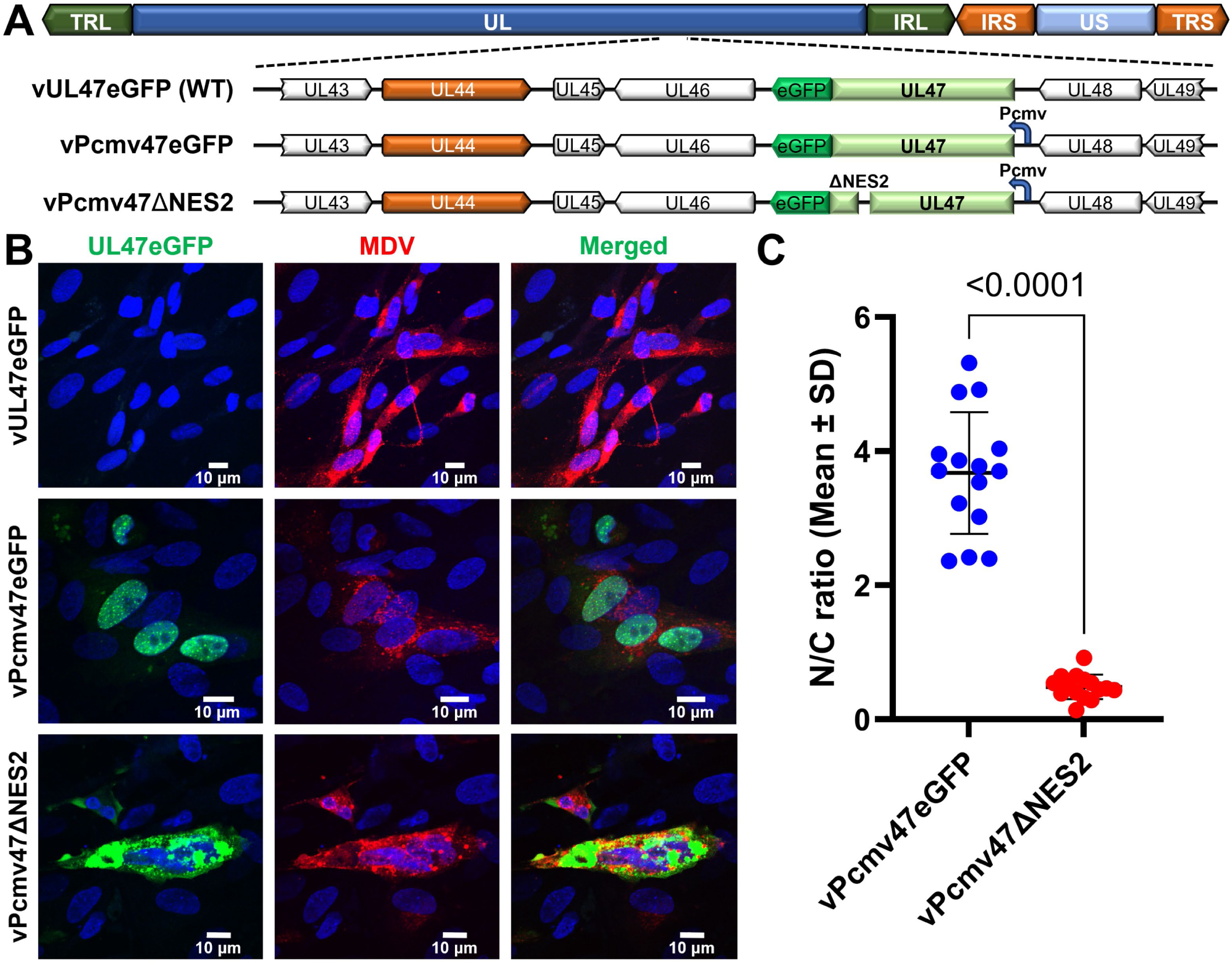
Subcellular localization of UL47eGFP and UL47eGFP lacking NES2 during infection. (A) Schematic representation of the MDV genome, with the UL47-encoding region amplified in the parental clone (vUL47eGFP), vPcmv47eGFP, and vPcmv47ΔNES2, generated by deleting the MDV UL47 NES2 motif (aa 664-678) in vPcmv47eGFP. (B) Representative images of UL47eGFP in CECs infected with vUL47eGFP, vPcmv47eGFP, and vPcmv47ΔNES2. Nuclei were stained with Hoechst 33342 (blue). Scale bars are 10 µm. (C) Mean nuclear-to-cytoplasmic ratios (N/C ratios) were computed, and individual quantified cells (n= 15) are shown in a scatter dot plot. UL47eGFP could not be readily detected in vUL47eGFP-infected CECs, so it was excluded from the analysis. We assessed data normality using the Shapiro-Wilk test. A two-tailed unpaired t-test with Welch’s correction was performed to assess differences between groups. Significant differences between the vPcmv47eGFP and vPcmv47ΔNES are indicated by p-values. Data are presented as the mean ± SD.

Next, we wanted to determine whether nucleocytoplasmic shuttling affected UL47_TEG5 function in replication and pathogenesis. Therefore, we created another deletion mutant where NES2 was removed (Δ664-678) in the wild-type MDV (vUL47eGFP) background to obtain (v47ΔNES2). We also generated another mutant in which the complete UL47_TEG5 ORF was removed, leaving eGFP (vΔ47). All recombinant viruses (Fig. 8A) were screened by RFLP analysis to confirm no extraneous changes occurred during the recombination process (Fig. S2). Following the reconstitution of vUL47eGFP, vΔ47, and v47ΔNES2 in cell culture, it was observed that eGFP intensity was strikingly higher in vΔ47-infected cells compared to vUL47eGFP and v47ΔNES2 (Fig. 8B). Measuring the relative eGFP intensity in infected cells confirmed that eGFP expression was significantly higher when not fused with UL47_TEG5 (Fig. 8C).

**Fig 8.**
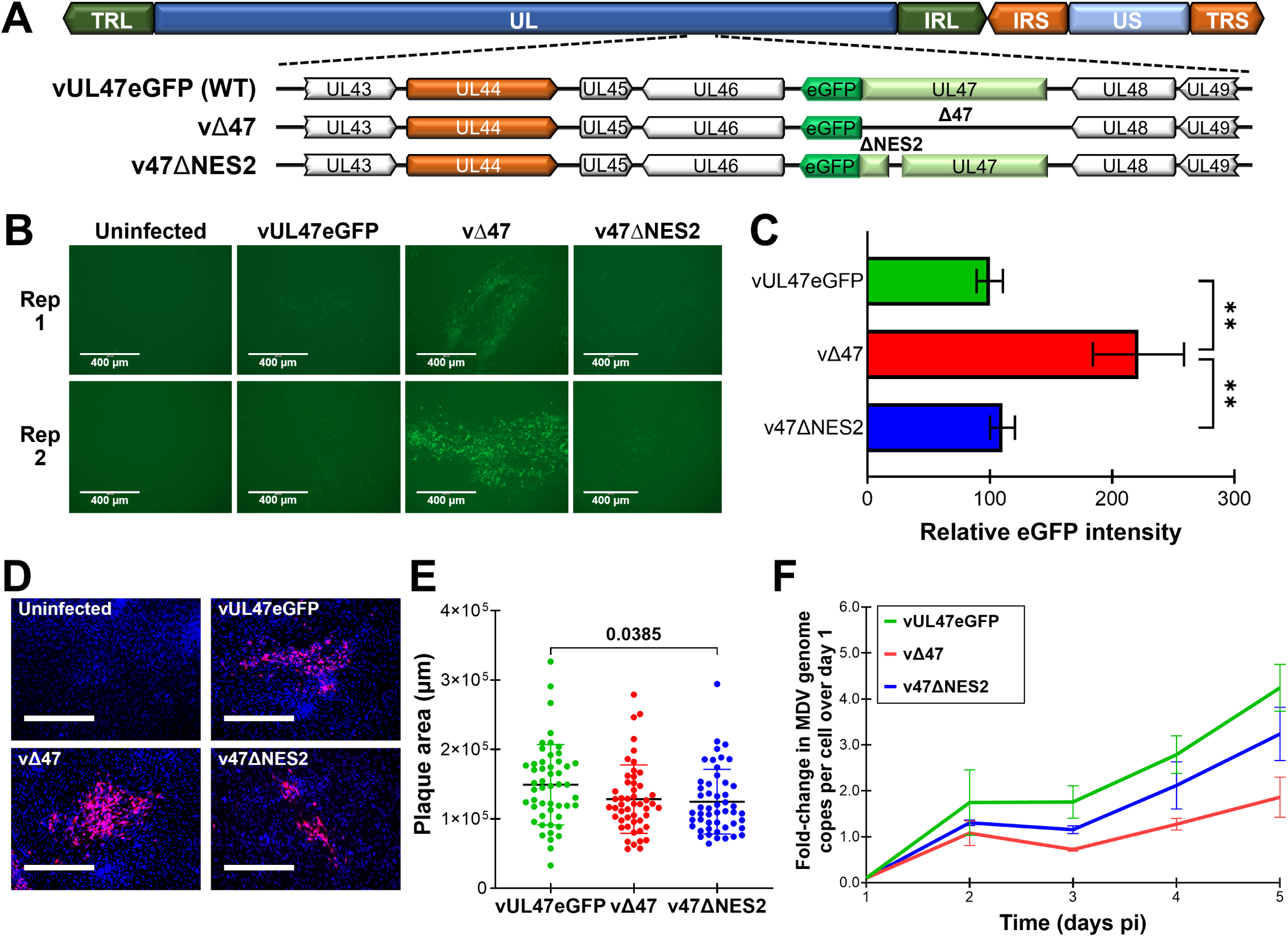
Characterization of rMDV in cell culture. (A) Schematic of the MDV BAC parental clone (vUL47eGFP) used to generate (vΔ47) and (v47ΔNES2). (B) Representative images showing direct eGFP imaging in vUL47eGFP-, vΔ47-, and v47ΔNES2-infected CECs. (C) Quantification of eGFP intensity expressed in plaques (n ≥ 10) produced by vUL47eGFP, vΔ47, and Δ47NES2 and expressed relative to vUL47eGFP. We assessed data normality using the Shapiro-Wilk test. We used a Kruskal-Wallis test followed by Dunn’s multiple-comparison test to assess differences between groups. Significant differences among vUL47eGFP, vΔ47, and v47ΔNES2 are indicated (**, p<0.01). Data are presented as the mean ± SD. (D) Representative plaques are shown in vUL47eGFP-, vΔ47-, and v47ΔNES2-infected CECs. (E) Plaque areas were measured from 50 individual plaques per virus at 4 days post-infection, and mean areas were computed and plotted as a scatter dot plot. We assessed data normality using the Shapiro-Wilk test. We used a Kruskal-Wallis test followed by Dunn’s multiple-comparison test to assess differences between groups. A significant difference between vUL47eGFP and v47ΔNES2 is indicated by the p-value. (F) Multistep growth kinetics were used to measure virus replication in CECs. Viral DNA was extracted daily from infected cells in triplicate and subjected to qPCR using gene-specific primers for the MDV ICP4 gene and the chicken iNOS gene as the internal control. We used the fold increase in viral DNA copies relative to day 1 to assess differences in replication. Data were assessed for normality using the Shapiro-Wilk test, and differences in viral replication between groups were evaluated using one-way ANOVA with Tukey’s multiple-comparison test. No significant difference was observed between groups.

To measure replication, plaque size and multi-step growth kinetic assays were used. There were no obvious differences in plaque sizes for all viruses (Fig. 8D), although plaque size assays showed significantly (p = 0.0385) smaller plaques induced by vΔ47NES2 (Fig. 8E). However, no differences were seen in multi-step growth kinetic assays between all three viruses (Fig. 8F). These result show deletion of UL47_TEG5 (vΔ47) and deletion of UL47_TEG5 NES2 (v47ΔNES2) had little effect on MDV replication in cell culture. In addition, these results suggest eGFP expression/intensity is severely reduced when fused to UL47_TEG5.

### Nucleocytoplasmic shuttling of MDV UL47_TEG5 is not required for replication or MD incidence in experimentally infected chickens

It has been previously shown that MDV UL47_TEG5 is dispensable for replication and oncogenicity in experimentally infected chickens but is required for horizontal transmission from chicken to chicken [17]. Here, we wanted to determine whether nucleocytoplasmic shuttling of MDV UL47_TEG5 is important for MDV replication, MD incidence (survival), and horizontal transmission. To do this, we experimentally infected chickens with vUL47eGFP (wild-type), vΔ47, or v47ΔNES2 and housed them with naïve contact chickens to measure transmission.

In experimentally infected chickens, viral genomes in the blood were quantified over the first five weeks to assess virus replication (Fig. 9A). At 28 days pi, v47ΔNES2 replication was significantly higher (p < 0.001) compared to vUL47eGFP and vΔ47; however, overall, there was no significance between each virus over the course of the experiment using two-way ANOVA. This was consistent with monitoring of feathers for viral infection, based on eGFP expression (Fig. 9C) and disease incidence (Fig. 9E), across the three viruses. These results show that deletion of UL47_TEG5, and more specifically UL47_TEG5 NES2 (aa 664-678), does not affect virus replication or disease induction in experimentally infected chickens.

**Fig 9.**
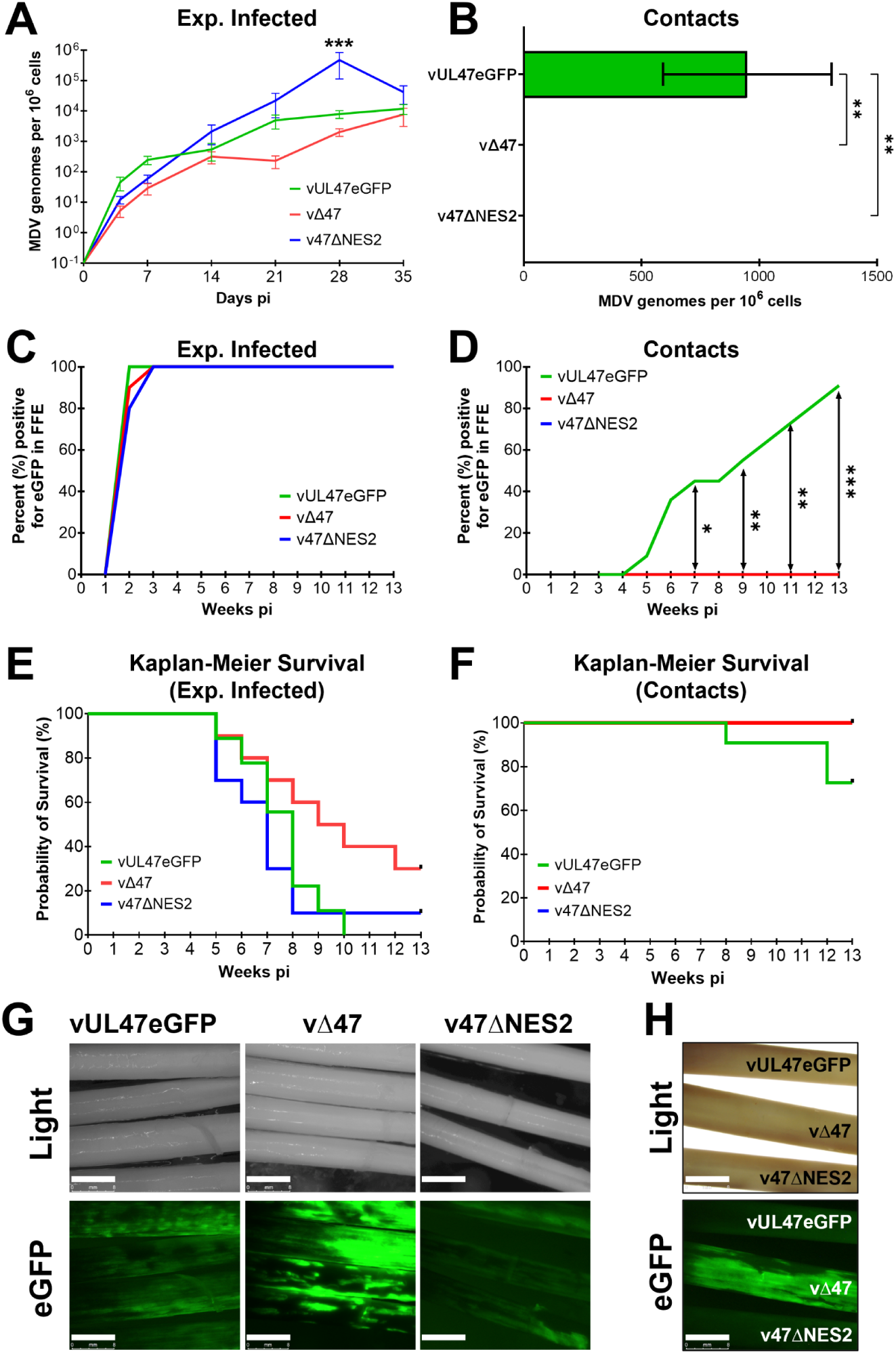
Replication and transmission of rMDVs in chickens. Chickens were experimentally infected with vUL47eGFP, vΔ47, or v47ΔNES2 as described in the Materials and Methods and monitored for 13 weeks. (A-B) We monitored replication in experimentally infected (A) and contact (B) chickens by quantifying MDV genomes in their blood over the first 5 weeks of infection (A) or at the termination of the experiment (B). We assessed data normality using the Shapiro-Wilk test. A two-way ANOVA followed by Tukey’s multiple-comparison test was performed to assess differences between groups. (A) A significant difference (***, p< 0.001) was observed only for v47ΔNES2 compared to vUL47eGFP and vΔ47 at 28 dpi. (B) No naïve contact chickens housed with vΔ47- or v47ΔNES2-infected chickens showed MDV genomes at termination, compared with contacts in the vUL47eGFP group. Shown are the mean MDV genomic copies per 106 blood cells ± standard error of the mean. (C-D) Quantitative analysis of the percent of birds positive for UL47eGFP or eGFP in feathers over the course of the experiment. (C) Using Fisher’s exact test at p < 0.05, there were no significant differences in the total number of chickens positive for experimentally infected chickens, with 100% positive by 21 days post-infection. (D) No naïve contact chickens housed with vΔ47- or v47ΔNES2-infected chickens were infected, while 91% of contact chickens were naturally infected with vUL47eGFP. Using Fisher’s exact test at p < 0.05, there was a significant difference between vUL47eGFP and the other two groups (vΔ47 and v47ΔNES2) between weeks 7 and 13. (E-F) Kaplan-Meier survival curves showing the probability of survival in experimentally infected (E) and contact (F) chickens over 13 weeks. Statistical comparisons between groups were performed using the Log-Rank (Mantel-Cox) test. (G) Representative feathers for each group are shown using direct fluorescent microscopy for UL47eGFP or eGFP at 35 days post-infection. Scale bars are 8 mm. (H) Representative feathers from each group for direct comparison.

To confirm that NES2 was important for the localization of MDV UL47_TEG5 in FFE cells, we examined the localization of UL47eGFP (vUL47eGFP and v47ΔNES or eGFP (vΔ47) in experimentally infected chickens using confocal microscopy. Single Z-stack images were used to examine the localization of eGFP with DNA in sections of FFs. Consistent with former reports [21, 22], UL47eGFP was nucleocytoplasmic in FFE cells (Fig. S3). In contrast, eGFP (vΔ47) and UL47eGFP lacking NES2 (vΔ47NES2) had a significantly negative correlation with DNA, supporting the strict cytoplasmic subcellular localization of UL47ΔNES2 or eGFP during infection with these viruses. These results show that MDV UL47_TEG5 NES2 is important for nucleocytoplasmic shuttling in FFE cells in chickens, but is not important for replication and MD in experimentally infected chickens.

### Nucleocytoplasmic shuttling of MDV UL47_TEG5 is required for transmission between chickens

To measure horizontal transmission, chickens housed with infected chickens were monitored for viral genomes in the blood (Fig. 9B), eGFP expression in feathers (Fig. 9D), and survival (Fig. 9F). Consistent with a previous report [17], vΔ47 failed to transmit, while vUL47eGFP efficiently transmitted to contact chickens, evidenced by viral genomes in the blood (Fig. 9B), eGFP expression in feather follicles (Fig. 9D), and MD induction (Fig. 9F). Interestingly, v47ΔNES2 also failed to infect contact chickens, as assessed by viral genomes in the blood (Fig. 9B), eGFP expression in feather follicles (Fig. 9D), and disease incidence (Fig. 9F). These results show that removal of NES2 from UL47_TEG5, rendering it unable to shuttle between the nucleus and cytoplasm (Fig. S3), is required for its functional requirement for horizontal transmission.

#### MDV gC expression and mRNA splicing of MDV UL44 (gC) in feather follicles

MDV gC is encoded by *UL44* and is required for horizontal transmission [19, 20]. MDV *UL44* mRNA is alternatively spliced to create variant products: membrane-bound MgC and two secreted forms, SgC104 and SgC145, all of which are required for efficient horizontal transmission [38].

Regulation of *UL44* mRNA splicing has been linked to both UL54_ICP27 and UL47_TEG5 [17]; however, more recent studies have questioned this [18]. We hypothesized that if MDV UL47_TEG5 regulated *UL44* mRNA splicing, preventing its nucleocytoplasmic shuttling by removing NES2 would affect *UL44* mRNA splicing. To further address this, we examined *UL44* mRNA splicing using RNA extracted from feather follicles of infected birds and performed RT-PCR assays to assess *UL44* mRNA products. As controls for mRNA expression, we also examined transcripts for MDV *UL54* (ICP27), MDV *UL48* (VP16), and cellular *GAPDH*. RT-PCR assays showed no detectable reduction in *UL44* (gC), *UL54*, or *UL48* mRNA levels in vΔ47- and v47ΔNES2-infected cells compared to vUL47eGFP (Fig. 10A). Importantly, we did not see any clear differences in *UL44* mRNA splicing between all three viruses, contrary to results previously published in vΔ47-infected cells [17]. Our results are consistent with a more recent report [18] that MDV UL47_TEG5 does not regulate *UL44* mRNA splicing.

**Fig 10.**
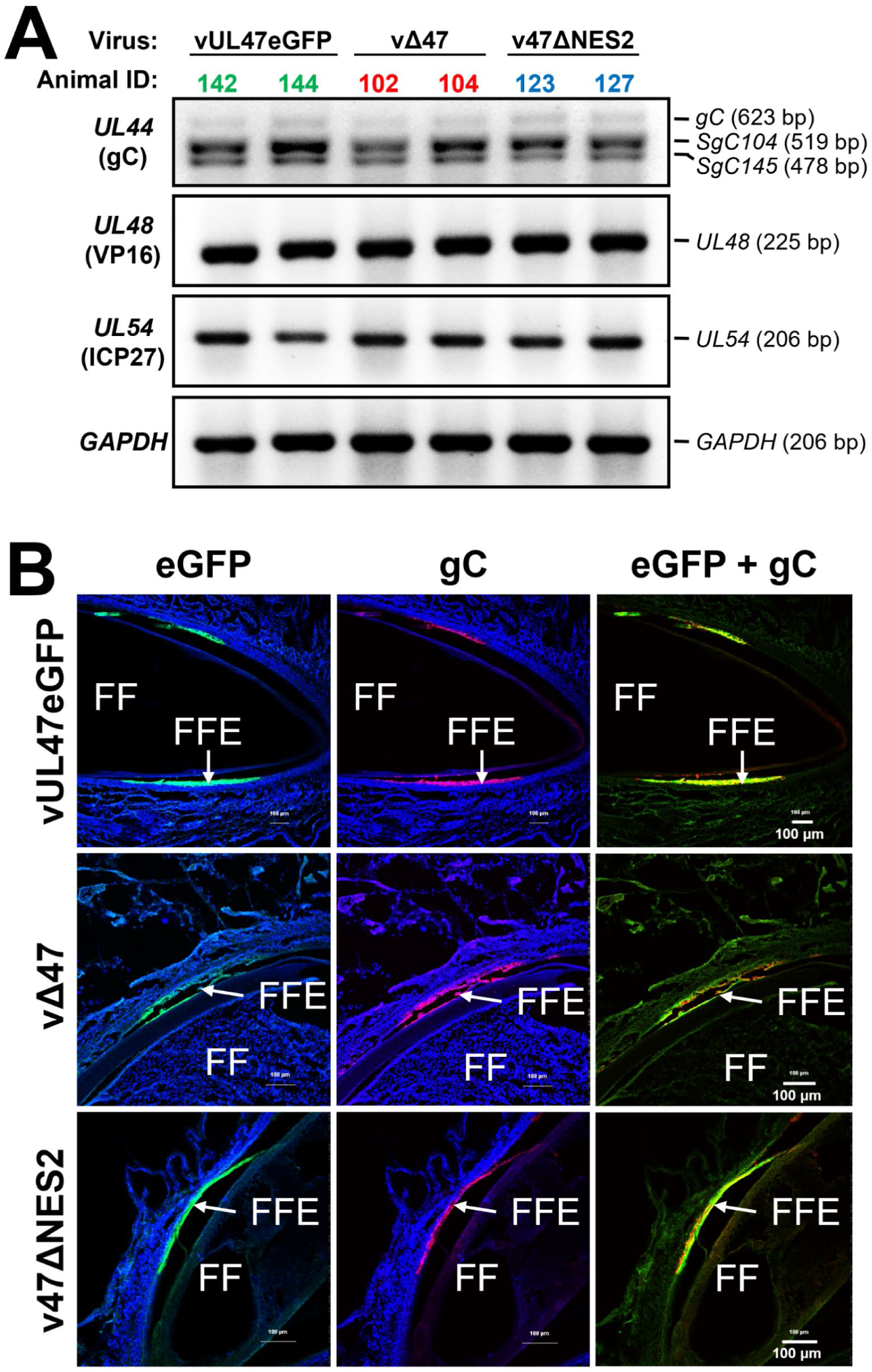
Expression of gC in FFE skin cells. (A) Total RNA was extracted from feathers of chickens experimentally infected with vUL47eGFP, vΔ47, or v47ΔNES2 and used in RT-PCR assays. We assessed UL44 (gC) splicing using primers spanning the splice region [62]. All three splice variants (gC-full, SgC104, SgC105) were expressed at their respective band sizes in three groups. We also used primers to detect *UL48* (VP16) and *UL54* (ICP27). *We used GAPDH* as a cellular control. (B) Representative images of skin/feather tissue cryosections cut transversely through the feather follicles (FF) infected with vUL47eGFP, vΔ47, or v47ΔNES2 at 28 days post-infection. FF and FF epithelial (FFE) skin cells are labeled in each image. Tissues were stained with mouse anti-gC A6 as the primary antibody. Anti-mouse IgM Alexa Fluor 568 was used as a secondary antibody. UL47eGFP (green) and Hoechst 33342 staining for DNA (blue) are also shown. The merged image shows eGFP and gC expression.

To further determine whether UL47_TEG5 affects gC expression outside of mRNA splicing, skin tissue collected from experimentally infected chickens was probed for gC using immunofluorescence assays and assessed by confocal microscopy. Using eGFP expression to mark infected cells, abundant gC was detected in skin from all three groups of infected chickens (Fig. 10B), indicating that neither UL47_TEG5 nor its nucleocytoplasmic shuttling is required for gC expression in FFE cells.

### Lack of localization of MDV UL47_TEG5 with chicken p32

It has also been reported that MDV UL47_TEG5 interacts with p32 within aa 71-185, and this interaction was important for horizontal transmission [18]. Although we did not delete the region believed to interact with p32/C1QBP, affecting this interaction could explain the lack of horizontal transmission of v47ΔNES2. Therefore, we examined colocalization of UL47_TEG5 with p32/C1QBP in cell culture. To do this, we first transfected DF-1 cells with pcUL47mRFP or pcUL47ΔNES2 and stained for p32/C1QBP antibody using immunofluorescence assays and confocal microscopy. Consistent with earlier results (Figs. 2-4), UL47_TEG5 was nuclear following transient transfection, while ΔNES2 was cytoplasmic (Fig. S4A). Endogenous p32/C1QBP was cytoplasmic and did not colocalize with UL47mRFP (nuclear), similar to other published work [18]. In contrast, UL47mRFP lacking NES2 (47ΔNES) was cytoplasmic and colocalized with p32/C1QBP in some areas. Next, we examined their localization during infection in cell culture. CECs infected with vPcmv47eGFP or Pcmv47ΔNES2 showed similar expression and p32/C1QPB staining (Fig. S4B) as in transient transfection. These results show that UL47_TEG5 and p32/C1QBP are not normally localized during transient transfection and cell culture infection, while deletion of NES2 renders UL47_TEG5 cytoplasmic, where it can colocalize with p32/C1QBP.

## DISCUSSION

MDV horizontal transmission or natural infection depends on fully productive replication in the FFE and shedding of infectious CFV [39]. Some viral proteins have been shown to be required for this process [19, 20, 31, 40], including UL47_TEG5 [17]; however, the mechanism by which UL47_TEG5 facilitates this process remains unclear. Previous studies showed that MDV UL47_TEG5 localizes primarily to the nucleus in cell culture, whereas it localizes predominantly to the cytoplasm in FFE skin cells (the site of CFV shedding) during the productive MDV replication cycle [21, 41]. This differential expression correlates with the cell-associated versus CFV stages of MDV replication, suggesting UL47_TEG5 subcellular trafficking may play a functional role in the assembly and release of enveloped CFVs. Since UL47_TEG5 is a conserved alphaherpesvirus tegument protein that shuttles between the nucleus and cytoplasm during infection [6], we sought to determine whether its ability to shuttle between the cytoplasm and nucleus was required for its function. Here, we mapped the NLS and NES motifs of MDV UL47_TEG5 and demonstrated that MDV UL47_TEG5 nucleocytoplasmic shuttling is critical for horizontal transmission in the natural host.

We used a systematic approach to identify regions within MDV UL47_TEG5 that are important for nucleocytoplasmic shuttling in cells. Deletion of individual putative NLS motifs did not alter the nuclear localization of UL47_TEG5 in transient transfection assays (Figs. 3 & 4). This was unexpected, but was consistent with what Durand *et al.* [18] observed when deleting MDV UL47_TEG5 aa 71-185 that contained the putative NLS1 (Fig. 1). NLS motifs have been defined for many alphaherpesviruses, including HSV-1, DEV, and BoHV-1, where N-terminal arginine-rich sequences are common [6, 8, 32, 42], much like MDV UL47_TEG5. Similarly, many contain C-terminal NES motifs [7, 10], again like MDV UL47_TEG5. HSV-1 UL47_TEG5 is the best characterized and contains a classical N-terminal NLS and a C-terminal CRM-1-dependent NES, allowing it to shuttle between the nucleus and cytoplasm [7, 8]. DEV UL47_TEG5 contains two NLS, one at the N-terminus (aa 40-50) and one at the C-terminus (aa 768-776) [42]. Our study extends this framework to MDV and suggests that nuclear import is not regulated by a single classical NLS. Deletion of putative NES motifs within the structured core shifted UL47_TEG5 localization toward the cytoplasm considerably. However, inhibition of CRM1/exportin-1 with LMB did not trap UL47_TEG5 NES mutants in the nucleus (Fig. 5). Verhagen *et al.* [10] identified a strong CRM1-independent NES at the N-terminus of BoHV-1 UL47_TEG5 (VP8). A broader, comprehensive analysis of HSV-1 tegument proteins revealed that nuclear export behavior can vary across proteins [43], supporting the idea that sequence-predicted export motifs may not function as traditional NES elements and emphasizing the need for experimental validation through localization assays.

The mechanism by which the removal of NES1 or NES2 alters the localization of MDV UL47_TEG5 is not fully understood, but misfolding is unlikely to be the cause. AlphaFold modeling identified a conserved UL47_TEG5 core (aa 400-800) that contained the NES motifs (Fig. 6B). Deletion of these NES motifs did not disrupt the predicted global protein folding (Fig. 6D), thereby arguing against misfolding as the basis of the localization phenotype and instead suggesting that the UL47_TEG5 core buried region contributes to nuclear retention or is required for regulated export.

Classical leucine-rich NES motifs typically mediate CRM1-dependent nuclear export, and deletion of such motifs often leads to strong nuclear accumulation. Removing an NES motif should trap the protein in the nucleus. Williams *et al.* [7] previously confirmed this for HSV-1 UL47_TEG5, in which the C-terminal NES functions via CRM1-mediated export and is sensitive to LMB, an inhibitor of CRM1-dependent nuclear export. Our data show that the subcellular localization of MDV UL47_TEG5 is not mediated by canonical mechanisms; it does not rely on CRM1-dependent export. Inhibition with LMB would be expected to result in enhanced nuclear accumulation of UL47_TEG5 ΔNES mutants, as observed for EIAV-Rev-GFP (Fig. 5). The absence of these effects suggests that the traditional import/export pathways do not play a dominant role in regulating the subcellular localization of UL47_TEG5. Instead, this supports a model in which UL47_TEG5 nucleocytoplasmic shuttling is regulated by a non-canonical mechanism or via protein-protein interactions rather than by classical NLS- or NES-mediated transport. Funk et al. [43] reported that LMB treatment resulted in HSV-1 UL47_TEG5 localizing exclusively to the nucleus and was insensitive to LMB treatment. They also noted that tegument proteins such as HSV-1 UL7_CEP1 and US2 showed nuclear export activity in their NEX-TRAP assay but lacked an obvious leucine-rich NES, suggesting they might still use a CRM-1-independent export mechanism. Verhagen *et al.* [9] used a GFP-tagged UL47-expressing recombinant virus to demonstrate that BoHV-1 UL47_TEG5 (VP8) localizes to the nucleus early in infection and undergoes rapid nucleocytoplasmic shuttling in infected cells. Further mapping revealed that BoHV-1 UL47_TEG5 (VP8) contains two NES motifs: a CRM1-dependent leucine-rich one and a novel CRM1-independent NES near the N-terminus (separable from the NLS). Full-length protein export is largely LMB-resistant due to the second NES. Together, these findings establish UL47_TEG5 nucleocytoplasmic shuttling as a conserved feature among alphaherpesviruses; however, its regulation may be virus-specific.

Using cell culture and experimental infection in chickens, we determined that nucleocytoplasmic shuttling of MDV UL47_TEG5 was not required for replication. In cell culture, the v47ΔNES mutants replicated comparably to the parental virus (Figs. 7 & 8). Similarly, when chickens were experimentally inoculated with the virus, there were no differences among the vUL47eGFP-, vΔ47-, and v47ΔNES2-infected groups (Fig. 9A, C, E), except that v47ΔNES2 replicated to higher levels at 28 days post-inoculation than vUL47eGFP and vΔ47. However, during natural infection, deletion of UL47_TEG5 (vΔ47) and UL47_TEG5 NES2 (v47ΔNES2) abrogated horizontal transmission in chickens (Fig. 9B, D, F), the former confirming previous reports by Chuard *et al.* [17]. Our results extend the requirement of UL47_TEG5 for horizontal transmission to its nucleocytoplasmic function.

In the previous report by Chuard *et al.* [17] it was suggested that UL47_TEG5’s role in facilitating horizontal transmission was through regulation of *UL44* (gC), specifically through mediating the splicing of *UL44* (gC). However, a more recent report by the same group suggested that its requirement for transmission may not be directly related to its regulation of gC, but rather to its interaction with p32 [18]. Our data here confirm UL47_TEG5 plays little to no role in *UL44* mRNA splicing and gC expression as evidenced by RT-PCR and immunoflouresence assays (IFA) of skin tissues that showed no defect in gC expression in vΔ47- and v47ΔNES2-infected skin tissues (Fig. 10). These results, in addition to the former studies, suggest UL47_TEG5 localization likely contributes to downstream processes, such as protein trafficking, virion maturation, and the release of infectious cell-free virions from FFE skin cells not directly related to gC.

Based on the abrogation of horizontal transmission when deleting the NES2 motif (aa 664-678) of MDV UL47_TEG5 (Fig. 9), mirroring the results of Durand *et al.* [18], we wondered whether deleting NES2 might also disrupt its reported interaction with p32/C1QBP previously mapped to aa 71-185. However, we observed no co-localization between MDV UL47_TEG5 and endogenous p32/C1QBP in either transient transfection or infection assays (Fig. S4). MDV UL47_TEG5 was strictly nuclear, while p32/C1QBP remained cytoplasmic—consistent with staining patterns reported by Durand *et al.* [18] using the same antibody. NES2 deletion shifted UL47_TEG5 to the cytoplasm, resulting in strong co-localization (Fig. S4). Thus, if the UL47_TEG5–p32/C1QBP interaction were critical for horizontal transmission, this cytoplasmic redistribution should have enhanced it. Instead, the opposite phenotype was observed, suggesting that the proposed interaction may not explain the transmission defect with v47ΔNES2 infection. Thus, the functional significance of the NES2 deletion mutant is likely not directly related to the requirement of p32 interaction with UL47_TEG5 in MDV horizontal transmission, but rather to other critical roles UL47_TEG5 plays in productive replication in FFE skin cells discussed below. However, further studies are needed to better define the UL47_TEG5 and nuclear p32 studied in Duran *et al.* [18].

Although individual NLS motif deletions had minimal impact on localization in cell culture, the triple NLS1-3 deletion caused only a modest shift (N/C ratio from ∼4 to ∼2; Fig. 3). Notably, UL47_TEG5 is mostly nuclear in cultured cells but nucleocytoplasmic in FFE skin cells (Fig. S3). This indicates that its localization is tightly regulated in relevant target cells, making cell-culture models suboptimal for studying these interactions. It would be valuable to test the functional importance of these putative NLS motifs during natural FFE skin cell infection and horizontal transmission, as they may play a greater role during natural infection.

Another key finding in our study is that fusing eGFP to UL47_TEG5 reduces eGFP expression in both cell culture and FFE skin cells (Figs. 8 & 9). In the vΔ47 mutant, eGFP levels were consistently and significantly higher in cell culture (Fig. 8B, C) and in FFE skin cells (Fig. 9G, H). This suggests that the UL47_TEG5 fusion destabilizes the protein, which may be something common among MDV proteins. We previously showed that several MDV proteins, including UL13_CHPK and US10, are minimally expressed in cell culture due to rapid degradation [31].

CHPK phosphorylates US10, thereby stabilizing it. Our recent data demonstrating CHPK-mediated phosphorylation of UL47_TEG5 [44], together with the low expression of the UL47eGFP fusion (Figs. 8 & 9), suggest that UL47_TEG5 may also be targeted for degradation and that CHPK phosphorylation may protect it from this fate. When eGFP is expressed alone, its levels are markedly higher than those of the UL47eGFP fusion. Further studies are needed to confirm this regulatory mechanism, but it suggests that MDV UL47_TEG5 may be unstable in infected cells, similar to UL13_CHPK and US10.

## SUMMARY

Our study links the nucleocytoplasmic shuttling of UL47_TEG5 to MDV horizontal transmission, likely through an essential role in regulating viral and cellular genes in FFE skin cells, thereby facilitating CFV production and dissemination. Figure 11 summarizes our working model of the requirement for MDV UL47_TEG5 for natural infection, likely by mediating CFV production in epithelial skin cells. Although data directly showing MDV UL47_TEG5 functioning in promoting fully productive CFV production are limited, we surmise it is a multifunctional protein important for multiple aspects of replication. MDV UL47_TEG5’s direct interaction with p32/C1QPB [18] suggests it plays a critical role in recruiting it to the nuclear membrane to help regulate the de-envelopment step during nuclear egress [45] or regulation of mRNA splicing [46]. MDV UL47_TEG5 interaction with and directing UL41_VHS to the cytoplasm has not been directly shown, but the inability of UL47ΔNES2 to shuttle to the nucleus would likely affect VHS function to selectively promote viral mRNA expression [47]. In addition, UL47_TEG augments UL46_VP16 activity in stimulating immediate-early gene expression [48]; thus, UL47ΔNES2 remaining in the cytoplasm would likely affect VP16 activity as well. It is important to note that cell-to-cell spread is unaffected by deletion or blocking the nucleocytoplasmic shuttling of UL47_TEG5 (47ΔNES2); therefore, its indispensable role in horizontal transmission is involved in the generation of CFV in FFE skin cells required for horizontal transmission. Alternatively, CFV may be produced but remain non-infectious due to a lack of proper tegumentation during virion assembly or a defect in tegument dissociation following entry into cells, as has been previously shown [49]. Although direct mechanistic studies are difficult using an *in vivo* model system like MDV in chickens, it provides a natural infection system to dissect cell-to-cell and cell-free virus spread in the natural host. It will be important to examine the specific hypotheses generated in this study, but our results show that UL47_TEG5’s ability to enter the nucleus is required for its critical role in CFV production and dissemination.

**Fig 11.**
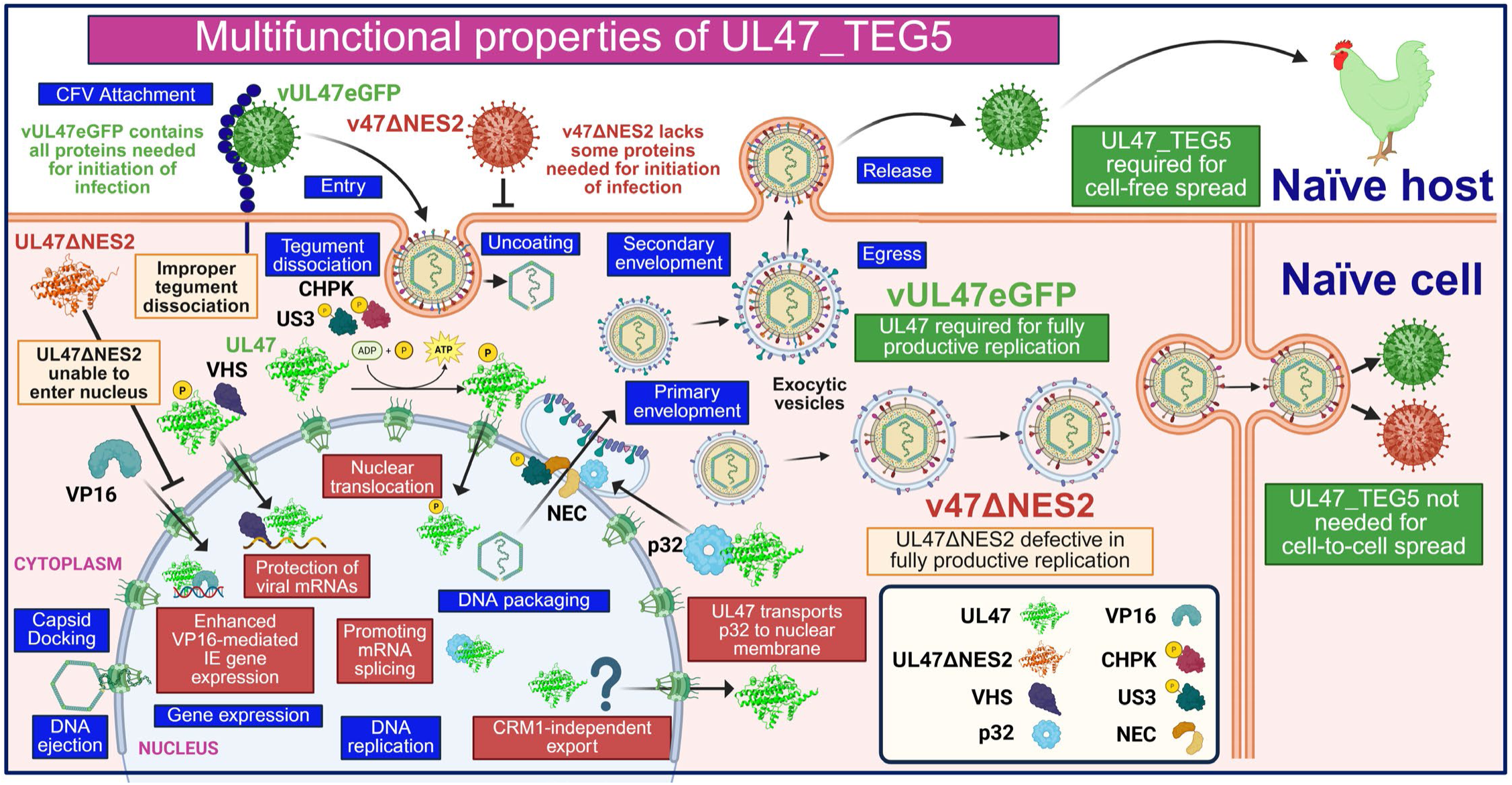
Working model for the multiple roles UL47_TEG5 may perform during horizontal transmission. UL47_TEG5 is required for CFV production and chicken-to-chicken spread (horizontal transmission), while dispensable for cell-to-cell spread. UL47_TEG5 homologs perform multiple functions during replication, including the promotion of tegument dissociation following virus entry, interaction and translocation of some UL41_VHS to the nucleus, where it protects viral mRNA from degradation, enhancing VP16-mediated transcription of immediate-early (IE) viral genes, and interaction with cellular p32/C1QBP that promotes mRNA splicing and nuclear egress. Phosphorylation of UL47_TEG5 by US3 or CHPK leads to nuclear import, whereas MDV UL47_TEG5 is exported from the nucleus through a CRM1-independent mechanism. Thus, UL47_TEG5 that cannot translocate to the nucleus (UL47ΔNES2) to perform these vital functions, leading to defective CFV production. However, UL47_TEG5 is not required for cell-to-cell spread. Figure created with BioRender.

## MATERIALS AND METHODS

### Cell culture and cells

Chicken embryo cells (CECs) were prepared using standard methods with 10–11-day-old specific-pathogen-free (SPF) eggs [50]. Primary CECs were cultured in Medium 199 (Cellgro, Corning, NY, USA) growth medium supplemented with 10% tryptose-phosphate broth (TPB), 0.63% NaHCO_3_ solution, antibiotics (100 U/mL penicillin and 100 µg/mL streptomycin), and 4% fetal bovine serum (FBS), and then changed to 0.2% FBS when the cells became confluent.

The immortalized chicken fibroblast cell line UNMSAH/DF-1 (ATCC CRL-12203) was grown in a 1:1 mixture of Leibovitz L-15 and McCoy 5A (LM) media (Gibco, Gaithersburg, MD, USA) supplemented with 10% FBS (LM10) plus antibiotics (100 U/mL penicillin and 100 µg/mL streptomycin). DF-1 cells stably expressing Cre recombinase (DF-1-Cre) were provided by Masahiro Niikura (USDA-ARS-ADOL, Lansing, MI) [51] and maintained in LM10 plus 50 µg/mL Zeocin (Invitrogen, Carlsbad, CA, USA).

All cells were maintained in a humidified atmosphere of 5% CO_2_ at 38 °C.

### Plasmids and transfections

The pcUL47mRFP expression plasmid was produced by cloning UL47mRFP from vUL47mRFP previously reported [30]. Briefly, UL47mRFP was PCR-amplified using Thermo Scientific Phusion Flash High-Fidelity PCR with the rUL47mRFP BAC clone as template and primers listed in S1 Table. The PCR product was resolved by agarose gel electrophoresis, then excised and purified using the Qiagen Gel Extraction Kit (Qiagen, Inc., Germantown, MD, USA). To produce pcUL47mRFP, the purified PCR product and the pcDNA3.1 vector (Thermo Fisher Scientific, Waltham, MA) were digested with BamHI and NotI, gel-purified, and then ligated using T4 DNA ligase (Invitrogen) for 16 h at 16°C. Ligation reactions were transformed into chemically competent *E. coli* DH5α, and transformants were selected on Luria-Bertani agar plates containing 100 µg/ml ampicillin. Plasmid DNA from positive clones was purified using the GeneJet Plasmid Miniprep Kit (Thermo Fisher Scientific). The final clone was Sanger sequenced at the University of Illinois DNA Services to confirm the absence of extraneous alterations.

Deletion mutants of pcUL47mRFP were generated by site-directed mutagenesis using the respective primers listed in S1 Table. Briefly, pcUL47mRFP was linearized using Thermo Scientific Phusion Flash High-Fidelity PCR master mix, and the PCR products were resolved by agarose gel electrophoresis, excised, and purified using the Qiagen Gel Extraction Kit (Qiagen), then ligated as described above. All mutations were verified by Sanger sequencing using primers flanking the targeted deletion regions and the *UL47* coding region. After generating pcUL47mRFP deletion mutants, expression plasmids were transfected into DF-1 cells using Lipofectamine 2000 reagent according to the manufacturer’s instructions.

### Antibodies

The following antibodies were used according to the manufacturers’ instructions or as previously published. For IFA with rMDVs in plaque size or subcellular localization assays, chicken anti-MDV polyclonal serum was used [52]. Goat anti-chicken IgY-Alexa Fluor 568 (Molecular Probes, Eugene, OR) was used as a secondary antibody. For IFA, mouse anti-gB MDV (IAN 86.17) [53], mouse anti-gC A6 [19], or anti-C1QBP/p32 (ProteinTech, catalog #24474-1-AP) primary antibodies were used. Anti-mouse IgG Alexa Fluor 568 (gB), anti-mouse IgM Alexa Fluor 568 (gC), and anti-rabbit IgG Alexa Fluor 488 or 568 were used as secondary antibodies. Hoechst 33342 (2 µg/ml, Molecular Probes) was used to visualize nuclei.

### Confocal microscopy and subcellular localization assays

DF-1 cells were seeded onto sterile glass coverslips in 6-well tissue culture dishes at ∼5 × 10^5^ cells per well and incubated overnight. The next day, cells (70-80% confluency) were transfected with 2.5 μg of plasmid DNA using Lipofectamine 2000 according to the manufacturer’s protocol. At 24-36 h post-transfection, cells were washed with phosphate-buffered saline (PBS) and fixed with 4% paraformaldehyde (PFA) in PBS buffer for 15 min at room temperature. Nuclei were stained with Hoechst 33342 (2 µg/ml, Molecular Probes). The fixed cells were mounted onto glass slides using ProLong Gold Antifade Mountant (Thermo Fisher Scientific) and incubated at room temperature overnight. Imaging was performed using a Nikon A1R laser-scanning confocal microscope with NIS-Elements C1 software. Images were captured with 40× or 60× oil-immersion objectives, using identical laser power, gain, and offset settings for comparisons.

### *In silico* prediction of NLS and NES sequences

Multiple *in silico* tools were used to identify putative NLS and NES motifs in MDV UL47_TEG5. Prediction of classical NLS motifs was performed using NLS Mapper [23], NLStradamus [24], NLSdb [25], seqNLS [26], and MyHits [27], which identify basic positively charged residues, while NES motifs were predicted using LocNES [28] that identifies leucine-rich sequences consistent with CRM1/exportin-1-dependent nuclear export. Motifs predicted with the highest confidence score and by at least two or more independent algorithms were selected for further analysis.

### Quantification of subcellular localization

Subcellular localization of UL47mRFP constructs was quantified using ImageJ/Fiji version 2.16.0 (https://imagej.net/ij/) for individual cells. For each cell, nuclear regions of interest (ROIs) were defined based on Hoechst 33342 staining, and cytoplasmic ROIs were calculated by subtracting the nuclear ROIs from the whole-cell ROIs. Mean fluorescent intensity values were measured for the nuclear and cytoplasmic compartments from the red channel (mRFP) or green channel (eGFP), and nuclear-to-cytoplasmic (N/C) ratios were calculated for each cell. For each construct, ≥12 cells per condition were analyzed [54, 55].

### Pharmacological inhibition of the nuclear transport pathway

At 24 h post-transfection, cells were treated with Leptomycin B (Sigma-Aldrich, Cat. No. L2913) at 20 nM in ethanol for 6 h to inhibit CRM1/exportin-1-dependent nuclear export. In parallel, cells were treated with ivermectin (Sigma-Aldrich, Cat. No. I8898) at 12.5 μM in DMSO for 1.5 h to inhibit importin α/β-mediated nuclear import. Vehicle controls included ethanol (EtOH) for LMB-treated cells and DMSO for IVM-treated cells, with volumes that matched those used in drug-treated cells. CRM1-dependent export control (EIAV-Rev-GFP) construct was included to verify inhibitor activity [56].

### 3D Protein modeling and geometry validation of UL47_TEG5

We used MDV UL47_TEG5 (UniProt: Q9E6M8) to predict the structure. 3D models of full-length UL47_TEG5 and ΔNES mutants were generated using AlphaFold2 implemented via ColabFold (ColabFold v1.6.1: AlphaFold2 using MMseqs2) under default settings [36]. The two putative NES motifs, NES1 (418-432) and NES2 (668-675), were removed *in silico* and submitted separately for structural prediction using identical parameters. We visualized predicted structures using UCSF ChimeraX version 1.10.1. Putative NLS and NES motifs were highlighted using residue-based coloring. We identified high-confidence domains using pLDDT scores and retained the core region (400-808) for comparative structural analysis. MDV UL47_TEG5, UL47_TEG5 ΔNES1, and UL47_ΔNES2 structures were superimposed in ChimeraX, and RMSD was calculated to assess any global change in protein folding from motif deletion. Structural geometry was assessed using MolProbity [57]. Ramachandran plots were used as a validation metric and to determine the proportion of residues in favored, allowed, and outlier regions.

### Recombinant (r) MDVs

The parental virus, vUL47eGFP, used here was previously reported [21], in which the eGFP coding sequence was inserted in frame at the C-terminus of MDV *UL47*. vPcmvUL47eGFP has also been published [22] and was generated by inserting the cytomegalovirus (CMV) immediate-early minimal promoter (Pcmv) upstream of *UL47eGFP* in vUL47eGFP. All BAC clones are based on the modified version of the RB-1B infectious BAC clone [58] restored for horizontal transmission [19].

The rΔUL47 mutant was made by deleting *UL47*, while rUL47ΔNES was generated by deleting the NES2 motif (664-678) in rUL47eGFP and rPcmvUL47eGFP. Briefly, the I-SceI-aphAI cassette was amplified from pEP-KanS2 by PCR using Thermo Fisher Scientific Phusion Flash High-Fidelity PCR Master Mix with primers listed in S2 Table. The PCR product was then used to mutagenize rUL47eGFP and rPcmvUL47eGFP in GS1783 *E. coli* cells using two-step Red recombination [59]. All BAC clones were screened by RFLP (Fig. S2), analytical PCR, and sequencing with primers for *UL47* previously described [21]. SnapGene 6.0.7 software (from Insightful Science; available at snapgene.com) was used to analyze gene sequencing results.

Viruses (v) were reconstituted by transfecting DF-1-Cre cells with purified BAC DNA using Lipofectamine 2000 (Invitrogen). After 2-3 days, transfected DF-1-Cre cells were mixed with freshly prepared primary CECs and further propagated in CECs until virus stocks were stored and titrated. All rMDVs were used at passage level four (p4) in all animal studies.

### Plaque size assays

Plaque areas were measured in CECs as previously described [52] using IFA. Briefly, primary anti-MDV chicken sera and secondary goat anti-chicken IgY-Alexa Fluor 568 secondary antibodies (Molecular Probes, Eugene, OR) were used to stain MDV plaques. Nuclei were counterstained with Hoechst 33342. For each rMDV, digital images of 50 individual plaques were collected using an EVOS FL Cell Imaging System (Thermo Fisher Scientific), and plaque areas were measured and quantified using ImageJ/Fiji version 2.16.0. Data distribution was evaluated for normality before analysis, and the Shapiro-Wilk test indicated that the normality assumptions were not met. A nonparametric analysis was performed using the Kruskal-Wallis test followed by Dunn’s multiple-comparison test to determine significant differences in GraphPad Prism version 10.6.1 (GraphPad Software). Scatter dot plots were generated, and a p-value of < 0.05 was considered statistically significant.

### Replication of rMDVs in cell culture

Multistep growth curves were used to assess viral replication in cell culture as previously described, using qPCR assays to measure viral DNA genomes [52]. Briefly, CECs were seeded in 6-well tissue culture plates and infected the following day with 100 plaque-forming units (PFU) of each virus per well. Total DNA was extracted from infected cells at 1, 2, 3, 4, and 5 days post-infection (pi) using DNA STAT60 from Tel-Test, Inc. (Friendship, TX, USA). Quantification of MDV genomic copies in CECs was performed using primers and probes to MDV ICP4 and chicken iNOS, as previously described [52]. Results were calculated as the viral-to-cellular genomes ratio. Each condition was analyzed in three technical replicates per qPCR reaction. The fold increase in viral DNA copies relative to day 1 was used to assess differences in replication. Data were assessed for normality using the Shapiro-Wilk test, and differences in viral replication between groups were evaluated using one-way ANOVA with Tukey’s multiple-comparison test. A p-value of <0.05 was considered statistically significant. All qPCR assays were performed on an Applied Biosystems QuantStudio 3 & 5 Real-Time PCR System (Thermo Fisher Scientific), and results were analyzed using the QuantStudio Design & Analysis Software v1.6.0 supplied by the manufacturer.

### RNA isolation and RT-PCR

Total RNA was collected using the RNA STAT-60 from Tel-Test, Inc. (Friendship, TX, USA) according to the manufacturer’s instructions. Briefly, vUL47eGFP-, vΔUL47-, and vUL47ΔNES-infected FFE skin cells were collected in RNA STAT60 as previously described [60]. Total RNA was DNase-treated using the Turbo DNA-*free* kit (Thermo Fisher Scientific) and stored at −80 °C until used for RT-PCR assays.

RT was performed with 1 µg of total RNA using the High-Capacity cDNA Reverse Transcription Kit (Thermo Fisher Scientific) with random priming, according to the manufacturer’s instructions. The reaction mixture was incubated at 25°C for 10 min, then at 37°C for 120 min, followed by 85°C for 5 min. To amplify cDNA, 3-5 µl of the RT mixture was mixed with DreamTaq Green PCR Master Mix (Thermo Fisher Scientific). Primers used for the amplification of the chicken *GAPDH*, *UL44*, *UL48*, and *UL54* have been previously published [22, 40]. For analysis of *UL44* (gC) splice variants, primers were used to amplify the expected transcript variants previously described [61]. All RT-PCR mixtures were electrophoresed through 2% agarose Tris-acetate-EDTA gels, and results were recorded using an AlphaImager HP imaging system (ProteinSimple, Minneapolis, MN, USA).

## Ethics statement

All animal work was conducted in accordance with national regulations. The animal care facilities and programs of the University of Illinois meet all the requirements of the law (89 –544, 91–579, 94 –276) and NIH regulations on laboratory animals and are in compliance with the Animal Welfare Act, PL 279, and are accredited by the Association for Assessment and Accreditation of Laboratory Animal Care (AAALAC). All experimental procedures were conducted in compliance with approved Institutional Animal Care and Use Committee protocols. Water and food were provided *ad libitum*.

### Animal experiments

SPF chickens were used for animal experiments. AVS Bio (Charles River Line 22) SPF eggs from non-MD vaccinated chickens were hatched and wing-banded at the University of Illinois Veterinary Medicine Research Farm (Urbana, IL), and then placed into three groups randomly (n=20/group). Five-day-old chicks were inoculated intra-abdominally with 2,000 plaque-forming units of vUL47eGFP, vΔ47, or v47ΔNES (n=10/virus) and housed with age-matched naïve contact chicks (n=10/group) to measure horizontal transmission (natural infection). To determine whether contact chickens were infected, UL47eGFP-negative birds were humanely euthanized at the termination of the experiment, and blood and serum were collected and analyzed by qPCR and IFA to confirm they were negative for MDV infection.

### Monitoring infection in feathers

To monitor the timing of when vUL47eGFP, vΔ47, and v47ΔNES reached the feathers, two flight feathers were plucked from both the right and left wings (totaling four feathers) of experimentally infected or contact-exposed chickens every week. Feathers were fixed in 2% PFA in PBS for 15 min, washed twice with PBS, and imaged directly using a Leica M205 FCA fluorescent stereomicroscope with a Leica DFC7000T digital color microscope camera (Leica Microsystems, Inc., Buffalo Grove, IL, USA). Some heavily infected chickens, based on intense fluorescent feathers, were euthanized the next day to collect feathers, skin sections, and other visceral organs for downstream studies.

### Viral growth kinetics *in vivo*

To measure virus replication in chickens, we collected whole blood at 4, 7, 14, 21, 28, and 35 days post-inoculation from experimentally infected birds. A total of 40 microliters of whole blood was collected by wing-vein puncture and mixed with 20 µl of 0.1 M EDTA. DNA was extracted using the E.Z. 96 Blood DNA Kit from Omega Bio-tek, Inc. (Norcross, GA), according to the manufacturer’s instructions. Primers and probes against MDV ICP4 and chicken iNOS were used in duplex qPCR reactions as previously described [52] in an Applied Biosystems QuantStudio 3 & 5 Real-Time PCR System (Thermo Fisher Scientific). The results were analyzed using QuantStudio Design & Analysis Software v1.6.0. The final viral loads were presented as MDV genomic copies per cell.

#### IFA of skin cryosections

MDV-infected skin/feather tissues were snap-frozen in Tissue Tek-optimal cutting temperature (OCT) compound (Sankura Finetek, Torrance, CA, USA) and stored at -80°C until sectioned. Eight- to ten-micrometer sections were affixed to Superfrost/Plus slides (Fisher Scientific, Pittsburgh, PA), fixed with 4% PFA in PBS buffer, washed twice with PBS, and then blocked with 10% fetal bovine serum (FBS) in PBS. Sections were incubated at 4°C overnight with mouse anti-gB MDV (IAN 86.17) [53], mouse anti-gC A6 [19], or anti-C1QBP/p32 (ProteinTech, catalog #24474-1-AP) primary antibodies diluted 1:200-500 in blocking buffer. Sections were washed three times with PBS, then incubated with anti-mouse IgG (gB), anti-mouse IgM (gC), or anti-rabbit IgG Alexa Fluor 568 or 488 secondary antibodies (1:500 in blocking buffer) for 1 h at room temperature. Nuclei were counterstained with Hoechst 33342 (2 µg/ml, Molecular Probes) for 5 min. Imaging was performed using a Nikon A1R laser-scanning confocal microscope with NIS-Elements C1 software. Images were captured using a 10× or 20× objective with identical laser power, gain, and offset settings.

## Statistical analysis

All statistical analyses were performed using GraphPad Prism version 10.6.1 (GraphPad Software). Data are presented as Mean ± SEM unless otherwise indicated. The statistical test used for each dataset is specified in the corresponding figure legend. Normality was assessed using the Shapiro-Wilk test. For normally distributed datasets, comparisons were made using one-way ANOVA with Tukey’s multiple-comparison test for more than two groups at a single time point. When normality was not met, the Kruskal-Wallis test with Dunn’s multiple-comparison test was used. For comparisons between two groups (N/C ratio for rMDVs), a two-tailed unpaired t-test with Welch’s correction was used. For comparisons among more than two groups over time or across different treatments, data were analyzed using a two-way ANOVA with Tukey’s multiple-comparison test. For survival analysis, Kaplan-Meier curves were compared using the Log-rank (Mantel-Cox) test. A p-value <0.05 was considered statistically significant.

## DATA AVAILABILITY STATEMENT

All relevant data are within the manuscript and its supporting information files.

## AUTHOR CONTRIBUTIONS

**Conceptualization:** Keith W. Jarosinski

**Data Curation:** Keith W. Jarosinski, Hafiz Sohaib Zafar

**Formal Analysis:** Keith W. Jarosinski, Hafiz Sohaib Zafar

**Funding Acquisition:** Keith W. Jarosinski

**Investigation:** Keith W. Jarosinski, Hafiz Sohaib Zafar

**Methodology:** Keith W. Jarosinski, Hafiz Sohaib Zafar

**Project Administration:** Keith W. Jarosinski

**Resources:** Keith W. Jarosinski, Hafiz Sohaib Zafar

**Supervision:** Keith W. Jarosinski

**Validation:** Keith W. Jarosinski, Hafiz Sohaib Zafar

**Visualization:** Keith W. Jarosinski, Hafiz Sohaib Zafar

**Writing—original draft preparation:** Hafiz Sohaib Zafar

**Writing—review and editing:** Keith W. Jarosinski, Hafiz Sohaib Zafar

Both authors have read and agreed to the published version of the manuscript.

## ACKNOWLEDGEMENTS

The authors thank Glorianna Wright, a DVM student in the College of Veterinary Medicine at Tuskegee University, for assistance with animal experiment sample collection during the Summer Research Training Program at the University of Illinois. This work used equipment, software, and facilities provided by the University of Illinois Urbana-Champaign College of Veterinary Medicine Shared Equipment Program.

## SUPPLEMENT INFORMATION CAPTIONS

**Fig S1. AlphaFold model quality assessment of UL47_TEG5 mutants.** (A) Representative top-ranked (rank_1) Predicted Aligned Error (PAE) maps generated with ColabFold for UL47_TEG5 and derived mutants. Blue regions indicate high confidence, whereas red regions indicate lower confidence in residue pair positions. (B) Ramachandran plots of the corresponding structures were generated using MolProbity for protein geometry analysis. Favored regions are highlighted, showing the percentage of residues in each region, and outlier residues are indicated for each model. The corresponding Ramachandran Z-score is also reported for each model.

**Fig S2. RFLP of rMDV BAC clones used in this report.** (A) Schematic representation of the MDV genome depicting the locations of the terminal repeat long (TRL) and short (TRS), internal repeat short (IRS), and unique long (UL) and short (US) regions. A portion of the UL region spanning *UL43*-*UL49* is expanded to better show changes in *UL47*. Differences between the rMDVs are shown. (B) Simulation and actual RFLP analysis of rMDV BAC clones. BAC DNA obtained for each parental (rUL47eGFP and rPcmv47eGFP), integrates (rΔ47-Int, r47ΔNES2-Int, and rPcmv47ΔNES2-Int), and resolved (rΔ47-Res, r47ΔNES2-Res, and rPcmv47ΔNES2-Res) BAC clones were digested with ClaI and examined using RFLP analysis. Insertion of the I-SceI-aphAI cassette during integration into the 17,667 bp fragment (red star) inserts a ClaI site, resulting in three fragments. Two fragments, 8,232 and 7,793, overlap (blue star), and a smaller fragment of 246 bp is not visible. Removal of the I-SceI-aphAI cassette during resolution yields a single 15,243-bp fragment (green star), in which *UL47* is removed. Deletion of the UL47 NES2 in rUL47eGFP results in three fragments: 10,221, 8,183, and 246 bp during integration (r47ΔNES2-Int), which resulted in a final fragment of 17,622 bp following resolution (r47ΔNES2-Res). Integration of the I-SceI-aphAI cassette into Pcmv47eGFP to delete UL47 NES (rPcmv47ΔNES2-Int) resulted in a shift of the 18,187 bp ClaI fragment into three (10,741, 8,183, and 246 bp) fragments (blue stars). Resolution restored the largest fragment from 18,187 to 18,142 bp (green star). Simulation was performed using SnapGene. The molecular weight marker used was the GeneRuler 1 kb Plus DNA Ladder from Thermo Scientific (Carlsbad, CA). No extraneous alterations are evident.

**Fig S3. Localization of eGFP in FFE skin cells.** (A) A representative Z-stack image of skin sections showing eGFP expression in FFE cells. Hoechst 33342 was used to identify cell DNA (nuclei). We used Z-stacks to generate linear ROIs for each group. Shown is a representative ROI used to measure the fluorescent intensity of eGFP and DNA (Hoechst 33342), along with their Pearson correlation coefficient (r). All confocal images used for quantitative analysis were captured on a Nikon confocal microscope with a 20× objective. We enabled the saturation lookup table (LUT) to identify and prevent saturated eGFP pixels, and used identical acquisition settings for all samples. (B) The Pearson correlation coefficient (r) between eGFP and Hoechst 33342 (DNA) was calculated for each ROI (n > 15) in each group. Pearson’s correlation coefficient (r) from individual ROIs was Fisher Z-transformed and averaged for each replicate for statistical comparison. Violin plots show the distribution of the correlation coefficient for each group. We assessed data for normality using the Shapiro-Wilk test, and evaluated differences among vUL47eGFP, vδ47, and v47ΔNES2 using a mixed-effects ANOVA. P-values indicate statistically significant differences. The average Pearson correlation coefficient (r) between eGFP and DNA was measured (vUL47eGFP = 0.6053 ± 0.1886; vΔ47 = 0.1389 ± 0.1753; v47ΔNES2 = 0.1167 ± 0.2116), showing that eGFP in FFE cells infected with vΔ47 or v47ΔNES2 was negatively correlated with DNA (nuclear) staining.

**Fig S4. Subcellular localization of UL47_TEG5 and cellular p32.** (A) DF-1 cells were transfected with pcDNA3.1 (EV), pcUL47mRFP (UL47), or pc47ΔNES2 for 24-36 h, fixed, and then stained with anti-p32 (green) or Hoechst 33342 (blue) to visualize nuclei. Shown is a representative image of >5 collected. (B) Representative images of CECs infected with recombinant vPcmvUL47eGFP, vPcmv47ΔNES2, or left uninfected. CECs were fixed after 5 days post-infection and stained with the anti-rabbit polyclonal C1QBP/p32 primary antibody (1:200), followed by the Alexa Fluor 568 anti-rabbit secondary antibody (1:500) for visualization (red). Nuclei were stained with Hoechst 33342 (blue).

**S1 Table. Primers used to generate mutations in expression plasmids.**

**S2 Table. Primers used to generate recombinant (r) Marek’s disease viruses (rMDV).**

