## Supplemental Figures for "Nucleocytoplasmic shuttling of the herpesvirus tegument protein UL47_TEG5 is required for its functional role in horizontal transmission in chickens"

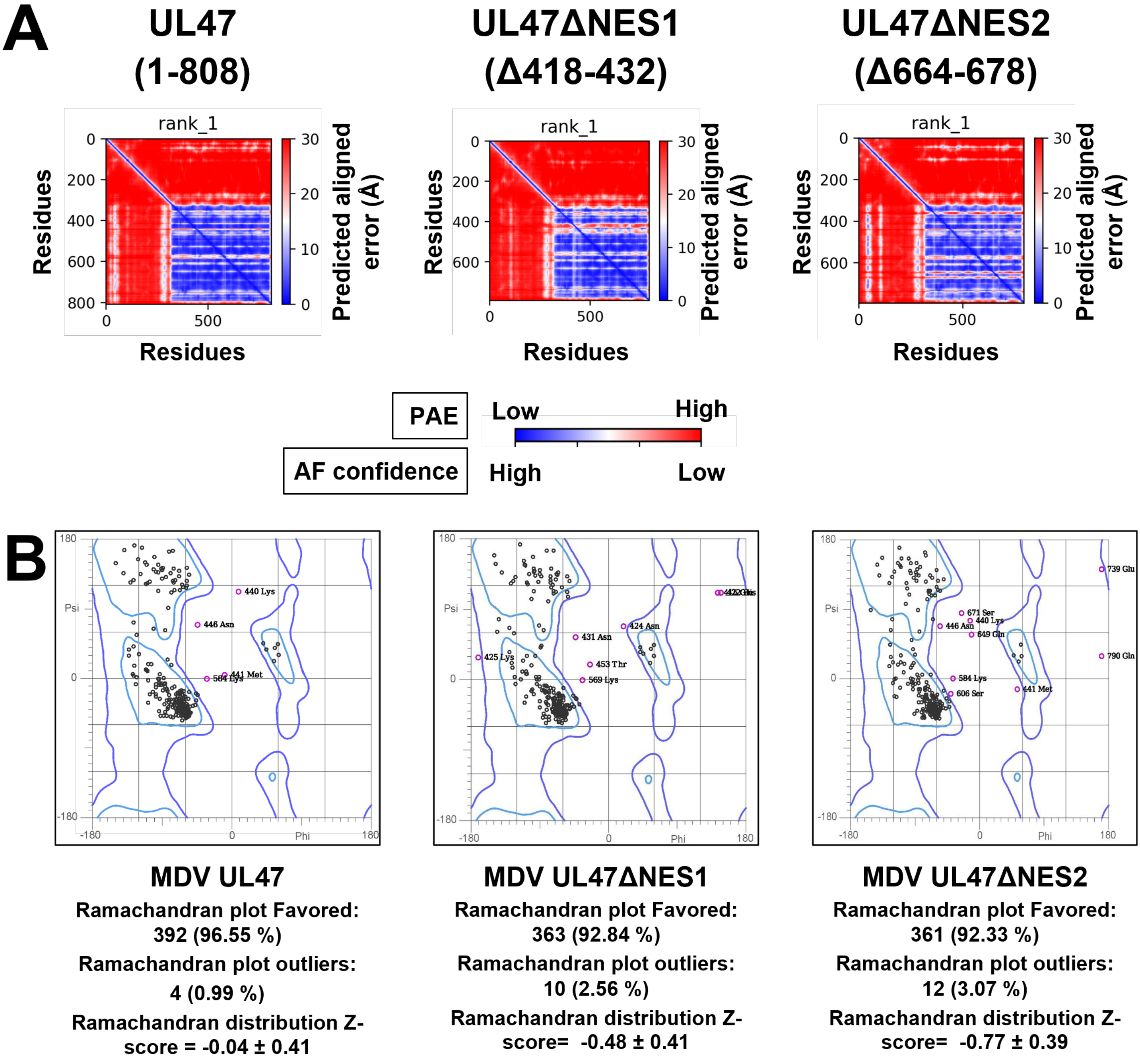


**Fig S1. AlphaFold model quality assessment of UL47_TEG5 mutants.** (A) Representative top-ranked (rank_1) Predicted Aligned Error (PAE) maps generated with ColabFold for UL47_TEG5 and derived mutants. Blue regions indicate high confidence, whereas red regions indicate lower confidence in residue pair positions. (B) Ramachandran plots of the corresponding structures were generated using MolProbity for protein geometry analysis. Favored regions are highlighted, showing the percentage of residues in each region, and outlier residues are indicated for each model. The corresponding Ramachandran Z-score is also reported for each model.

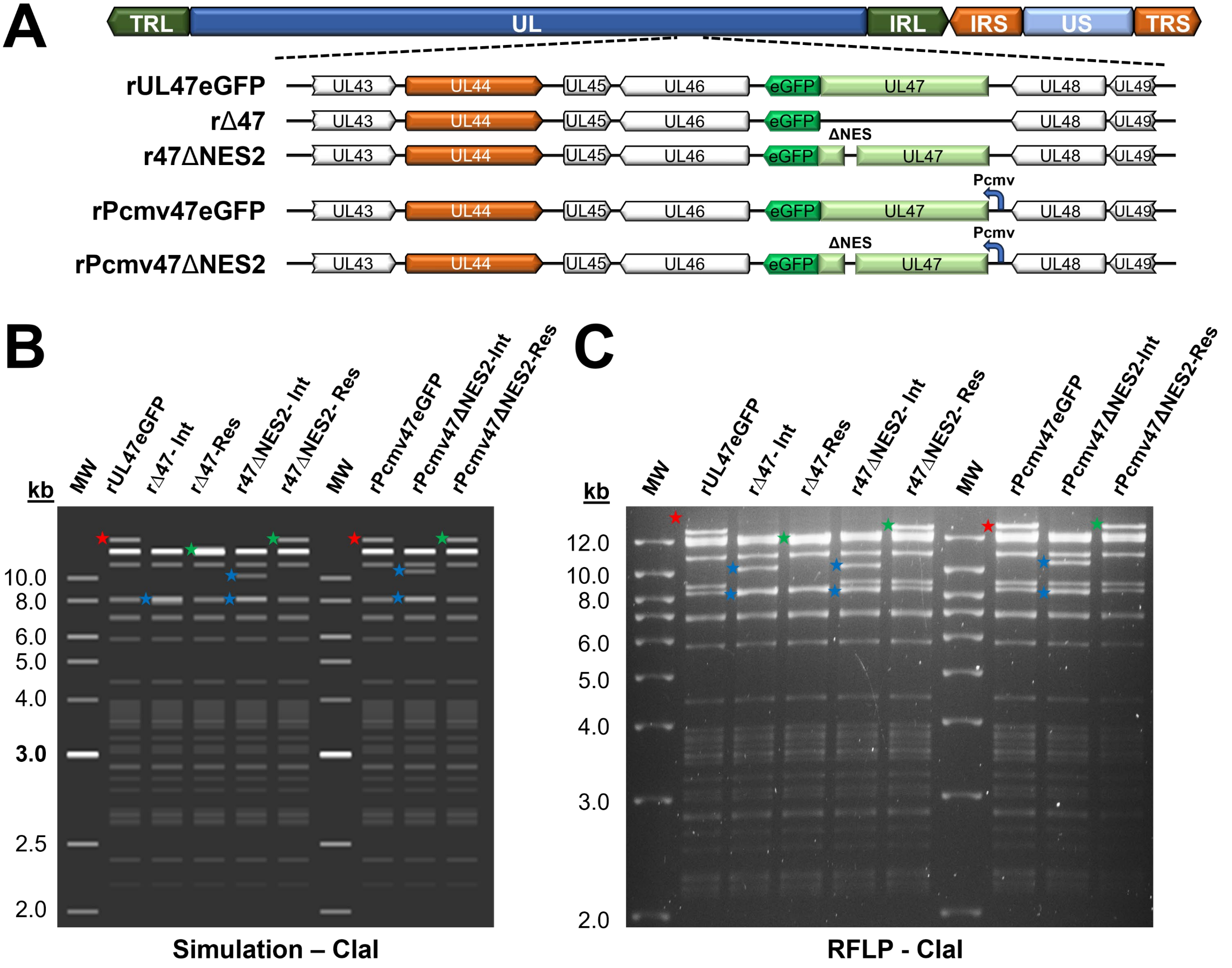


**Fig S2. RFLP of rMDV BAC clones used in this report.** (A) Schematic representation of the MDV genome depicting the locations of the terminal repeat long (TRL) and short (TRS), internal repeat short (IRS), and unique long (UL) and short (US) regions. A portion of the UL region spanning UL43-UL49 is expanded to better show changes in UL47. Differences between the rMDVs are shown. (B) Simulation and actual RFLP analysis of rMDV BAC clones. BAC DNA obtained for each parental (rUL47eGFP and rPcmv47eGFP), integrates (rΔ47-Int, r47ΔNES2-Int, and rPcmv47ΔNES2-Int), and resolved (rΔ47-Res, r47ΔNES2-Res, and rPcmv47ΔNES2-Res) BAC clones were digested with ClaI and examined using RFLP analysis. Insertion of the I-SceI-aphAI cassette during integration into the 17,667 bp fragment (red star) inserts a ClaI site, resulting in three fragments. Two fragments, 8,232 and 7,793, overlap (blue star), and a smaller fragment of 246 bp is not visible. Removal of the I-SceI-aphAI cassette during resolution yields a single 15,243-bp fragment (green start), in which UL47 is removed. Deletion of the UL47 NES2 in rUL47eGFP results in three fragments: 10,221, 8,183, and 246 bp during integration (r47ΔNES2-Int), which resulted in a final fragment of 17,622 bp following resolution (r47ΔNES2-Res). Integration of the I-SceI-aphAI cassette into Pcmv47eGFP to delete UL47 NES (rPcmv47ΔNES2-Int) resulted in a shift of the 18,187 bp ClaI fragment into three (10,741, 8,183, and 246 bp) fragments (blue stars). Resolution restored the largest fragment from 18,187 to 18,142 bp (green star). Simulation was performed using SnapGene. The molecular weight marker used was the GeneRuler 1 kb Plus DNA Ladder from Thermo Scientific (Carlsbad, CA). No extraneous alterations are evident.

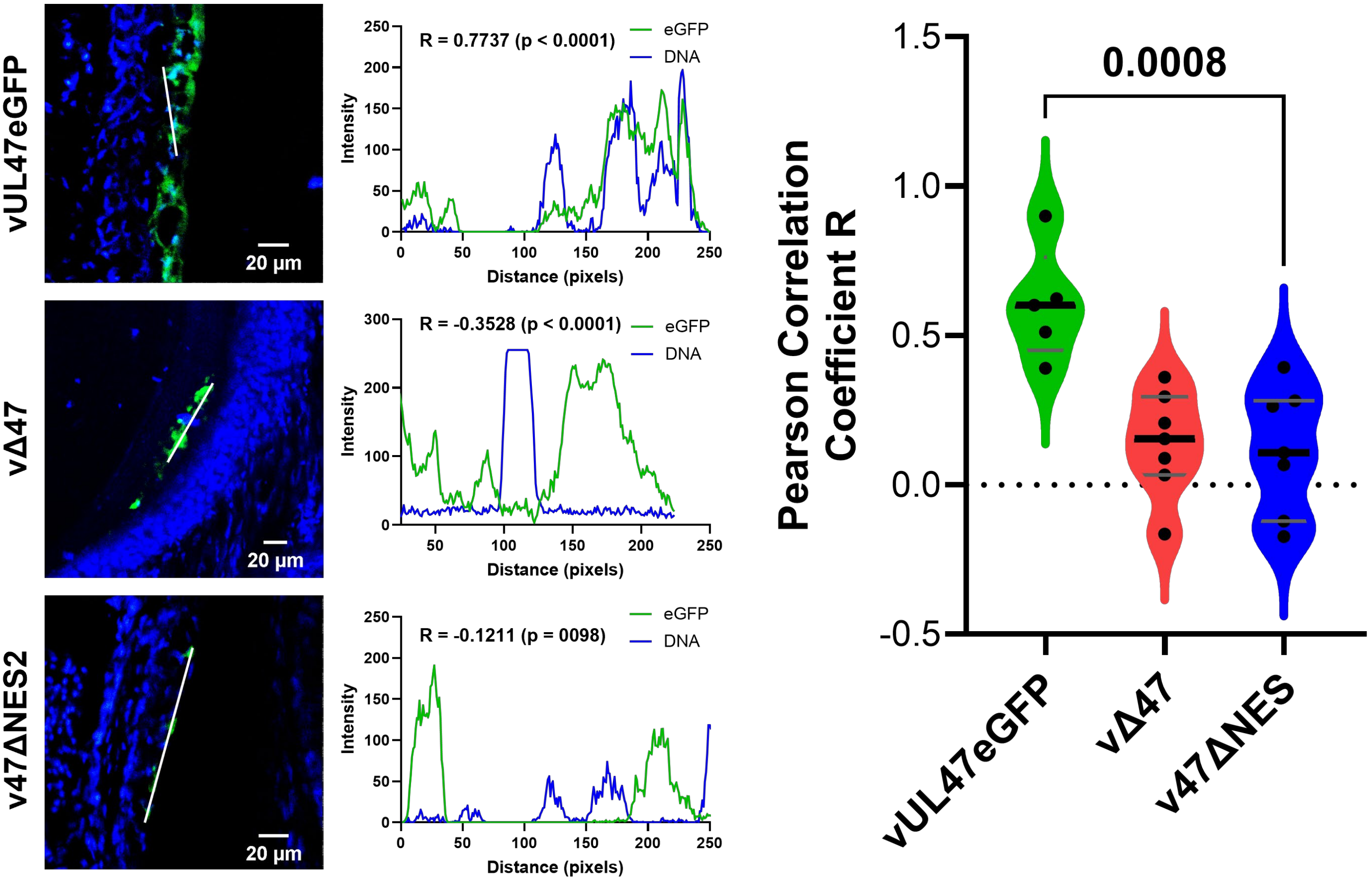


**Fig S3. Localization of eGFP in FFE skin cells.** (A) A representative Z-stack image of skin sections showing eGFP expression in FFE cells. Hoechst 33342 was used to identify cell DNA (nuclei). We used Z-stacks to generate linear ROIs for each group. Shown is a representative ROI used to measure the fluorescent intensity of eGFP and DNA (Hoechst 33342), along with their Pearson correlation coefficient (r). All confocal images used for quantitative analysis were captured on a Nikon confocal microscope with a 20× objective. We enabled the saturation lookup table (LUT) to identify and prevent saturated eGFP pixels, and used identical acquisition settings for all samples. (B) The Pearson correlation coefficient (r) between eGFP and Hoechst 33342 (DNA) was calculated for each ROI (n > 15) in each group. Pearson's correlation coefficient (r) from individual ROIs was Fisher Z-transformed and averaged for each replicate for statistical comparison. Violin plots show the distribution of the correlation coefficient for each group. We assessed data for normality using the Shapiro-Wilk test, and evaluated differences among vUL47eGFP, vδ47, and v47ΔNES2 using a mixed-effects ANOVA. P-values indicate statistically significant differences. The average Pearson correlation coefficient (r) between eGFP and DNA was measured (vUL47eGFP = 0.6053 ± 0.1886; vΔ47 = 0.1389 ± 0.1753; v47ΔNES2 = 0.1167 ± 0.2116), showing that eGFP in FFE cells infected with vΔ47 or v47ΔNES2 was negatively correlated with DNA (nuclear) staining.

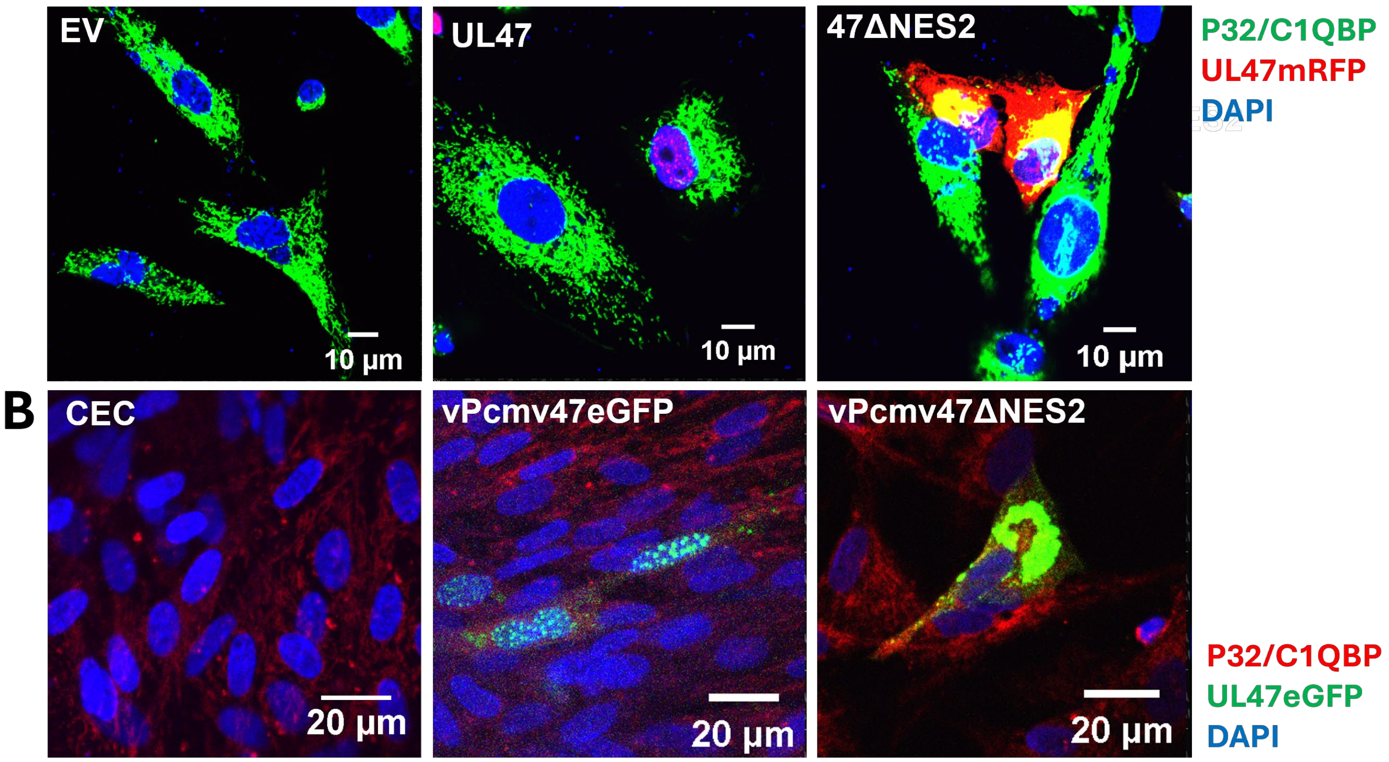


**Fig S4. Subcellular localization of UL47_TEG5 and p32.** (A) DF-1 cells were transfected with pcDNA3.1 (EV), pcUL47mRFP (UL47), or pc47ΔNES2 for 24-36 h, fixed 24-36 h post-transfection, and stained with anti-p32 (green) or Hoechst 33342 (blue) to visualize nuclei. Shown is a representative image of >5 collected. (B) Representative images of CECs infected with recombinant vPcmvUL47eGFP, vPcmv47ΔNES2 or left uninfected. CECs were fixed after 5 days pi and stained with the anti-rabbit polyclonal C1QBP/p32 primary antibody (1:200), followed by the Alexa Fluor 568 anti-rabbit secondary antibody (1:500) for visualization (red). Nuclei were stained with Hoechst (blue).
