## Supplemental Tables for "Nucleocytoplasmic shuttling of the herpesvirus tegument protein UL47_TEG5 is required for its functional role in horizontal transmission in chickens"

**S1 Table. Primers used to generate mutations in expression plasmids.**

| **Construct[a]** | **Primer Name[b]** | **Sequence (5'- 3')** |
|---|---|---|
| pcUL47mRFP[c] | UL47ForBam | TATTGGATCCTATGCTGGTAGCACATTCCACCG |
| | UL47RevNot | TTATGAGCGGCCGCTGCTGCAAGAGTCCCGGTTTTAA |
| Δ2-149 | DEL_NLS1_UL47For | GCCAGCATGCATATGCATTTTC |
| | DEL_seq 2-149_UL47Rev | CATTCTGGATCCGAGCTCGGTA |
| Δ150-300 | DEL_NLS3_UL47For | ACTGGTGATAGTAAAGGTACGAGTC |
| | DEL_seq 150-300_UL47Rev | ATTACGATGGCGTCGTCTAGATG |
| Δ301-450 | DEL_seq 301-450_UL47For | TCCACAATGCCACACGTTCG |
| | DEL_seq 301-450_UL47Rev | GGTACGCTTATAGGTTTTTAGCGACT |
| 7Δ451-600 | DEL_seq 451-600_UL47For | CTTGGGAGGTTAGAGTCATCTCT |
| | DEL_seq 451-600_UL47Rev | TAATATTTTTACATTTCCATATTTATGCATTTTATTTTTATGT |
| Δ601-808 | DEL_seq 601-808_UL47For | ATGGCCTCCTCCGAGGAC |
| | DEL_seq 601-808_UL47Rev | GCAGGCCCCCTCTGCATAA |
| ΔNLS1 | DEL_NLS1_UL47For | GCCAGCATGCATATGCATTTTC |
| | DEL_NLS1_UL47Rev | TAATGATGCATTAATCATATCACGAGATG |
| ΔNLS2 | DEL_NLS2_UL47For | GGTCACGGAGATTCTTTTACATCC |
| | DEL_NLS2_UL47Rev | TGATGCATGATTATATTCTCCTTCTCT |
| ΔNLS3 | DEL_NLS3_UL47For | ACTGGTGATAGTAAAGGTACGAGTC |
| | DEL_NLS3_UL47Rev | GTAATTGTCTTCTCCAAAACTATCATCTAT |
| ΔNES1 | DEL_NES2_UL47For | GCAATTTGTATACATAAAAATAAAATGCATAAATA |
| | DEL_NES2_UL47Rev | ATTCAACTCACGCAATGACAT |
| ΔNES2 | DEL_NES_UL47For | ATCGCGCAGATGGTTATTGG |
| | DEL_NES_UL47Rev | AGCAATCACGGCAGCG |

| | | |
|---|---|---|
| ΔNLS1+2[d] | DEL_NLS2_UL47For<br>DEL_NLS2_UL47Rev | GGTCACGGAGATTCTTTTACATCC<br>TGATGCATGATTATATTCTCCTTCTCT |
| ΔNLS1+3[e] | DEL_NLS1_UL47For<br>DEL_NLS1_UL47Rev | GCCAGCATGCATATGCATTTTC<br>TAATGATGCATTAATCATATCACGAGATG |
| ΔNLS2+3[e] | DEL_NLS2_UL47For<br>DEL_NLS2_UL47Rev | GGTCACGGAGATTCTTTTACATCC<br>TGATGCATGATTATATTCTCCTTCTCT |
| ΔNLS1+2+3[f] | DEL_NLS3_UL47For<br>DEL_NLS3_UL47Rev | ACTGGTGATAGTAAAGGTACGAGTC<br>GTAATTGTCTTCTCCAAAACTATCATCTAT |

[a]Expression construct generated and used as a template for mutations unless noted.
[b]Name of the primers.
[c]Restriction digestion cloning using BamHI and NotI restriction enzymes (underlined) to amplify UL47mRFP from vUL47mRFP and clone it into the pcDNA3.1 cloning vector.
[d]ΔNLS1 as template
[e]ΔNLS3 as template
[f]ΔNLS1+2 as template

**S2 Table. Primers used to generate recombinant (r) Marek's disease viruses (rMDV).**

| Modification[a] | Primer Name | Sequence (5'- 3')[b] |
|---|---|---|
| rΔ47 | EP_DeltaMDVUL47for | GTTTTGATAACAGTATGCTGGTAGCACATTCCACCGAAGAATGGTGAGCAAGGGCGAGGA*TAGGGATAACAGGGTAATCGATTT* |
| | EP_DeltaMDVUL47rev | GCACCACCCCGGTGAACAGCTCCTCGCCCTTGCTCACCATTCTTCGGTGGAATGTGCTAC*GCCAGTGTTACAACCAATTAACC* |
| r47ΔNES2; rPcmv47ΔNES2 | EP_DeltaMDVUL47_NESfor | TGTCAAGCTGGGAGAAAAATTAACCGCTGCCGTGATTGCTATCGCGCAGATGGTTATTGG*TAGGGATAACAGGGTAATCGATTT* |
| | EP_DeltaMDVUL47_NESrev | CTTTTTTGTGATATACAGATCCAATAACCATCTGCGCGATAGCAATCACGGCAGCGGTTA*GCCAGTGTTACAACCAATTAACC* |

[a]Modification produced in the recombinant virus, including deletion of *UL47* (ΔUL47) and the NES motif (aa 668-678) from UL47, leaving behind eGFP

[b]Red indicates unique upstream integration sequences. Green indicates unique downstream integration sequences. Blue indicates complementary sequences used to resolve integrations. *Italics indicate the template-binding region of the primers for PCR amplification with pEP-KanS2.*
